# Fluidity and Predictability of Epistasis on an Intragenic Fitness Landscape

**DOI:** 10.1101/2024.08.25.609583

**Authors:** Sarvesh Baheti, Namratha Raj, Supreet Saini

## Abstract

How epistasis hinders or facilitates movement on fitness landscapes has been a longstanding question in evolutionary biology. High-throughput experiments have revealed that, despite their idiosyncratic nature, epistatic interactions can exhibit reproducible global statistical patterns. Recently, Papkou *et al.* constructed a fitness landscape for a 9-base pair region of the *folA* gene in *Escherichia coli*, which encodes dihydrofolate reductase (DHFR), and showed that this landscape is both rugged and highly navigable. Here, we analyze this landscape to address two questions: How does the nature of epistasis between two mutations change with genetic background? and How predictable is epistasis within a gene? We find that epistasis is “fluid”: higher-order interactions cause the relationship between two mutations to shift strongly across genetic backgrounds. Mutations fall into two distinct categories: a small subset exhibit strong global epistasis, while the majority do not. Nonetheless, we find that the distribution of fitness effects (DFE) of a genotype is highly predictable from its fitness. These findings provide a gene-level perspective on how epistasis operates, revealing both its unpredictability and its statistical regularities, and offer a framework for predicting mutational effects from high-dimensional fitness landscapes.

**Significance Statement.:** The effect of a mutation on fitness depends on the genome in which it occurs, a phenomenon known as epistasis. Epistasis makes evolution difficult to predict, but recent work has uncovered statistical regularities in how it manifests. Using a fitness landscape of ∼260,000 variants of an *E. coli* gene, we show that higher-order interactions make pair-wise epistasis “fluid”: the relationship between two mutations changes with genetic background. We also find that epistasis is “binary” - a small subset of mutations exhibits strong global statistical patterns, while most do not. These findings reveal new principles of how epistasis shapes protein evolution and, ultimately, organismal adaptation.

## Introduction

Mutations determine the fitness of an organism in a genetic background-dependent manner, a phenomenon known as epistasis^1^. Epistasis shapes adaptation^2,3^ and plays a central role in evolutionary dynamics, but it remains largely unpredictable^4–6^. In recent years, however, a few reproducible statistical patterns of epistasis, often linked to macroscopic traits, have begun to emerge^7–13^. One way to study such patterns is through the concept of the fitness landscape^14–16^.

A fitness landscape can be visualized as a multidimensional surface, where one axis represents fitness and the remaining axes represent genotype. It thus provides a genotype–phenotype map whose shape, in a given environment, is dictated by genetic interactions and epistasis^11,17–19^. Landscapes with many local peaks are termed rugged^20–22^, and on such landscapes, populations may become trapped at suboptimal fitness peaks, with outcomes strongly dependent on chance events and initial genetic states^23–25^. By contrast, smooth landscapes allow populations starting from diverse points in sequence space to converge on the same global optimum^26–29^. Therefore, the topography of a fitness landscape directly influences the predictability of evolution, with broad implications for both theory and application.

Although fitness landscapes were originally conceived as abstract models linking genotype to fitness, they have become particularly powerful tools to study protein sequence–function-fitness relationships^30^. Early efforts focused on landscapes defined by a small number of “key” sites, typically treated as biallelic^29,31–39^. While these approaches revealed the ubiquity of epistasis in proteins, their limited dimensionality restricted the scope for statistical generalization^40^. Specifically, how does the nature of intramolecular epistasis scale beyond a handful of sites? For example, positive epistasis in β-lactamases has been shown to enhance catalytic efficiency and facilitate the evolution of antibiotic resistance, with broad implications for host–pathogen interactions and drug resistance evolution^41^. The advent of high-throughput mutagenesis and phenotyping technologies has since enabled the construction of high-dimensional landscapes, capturing thousands of genotypes and making systematic exploration of epistasis possible^42–48^.

In this study, we analyze one such high-dimensional landscape, constructed by Papkou and coworkers^43^, that captures the fitness of all possible sequences in a 9–base pair region of the *folA* gene in *Escherichia coli* (*E. coli*). *folA* encodes dihydrofolate reductase (DHFR), an essential enzyme whose mutations are well known to confer resistance to the antibiotic trimethoprim^49–53^.

We use this landscape to address three central questions (**Supplement Figure S1**): How does the nature of epistasis between two sites change as a function of genetic background? Are these changes determined primarily by genotype or by background fitness? Do mutations follow strong patterns of global epistasis? And if so, what governs this behavior?

Through our analyses, we show that epistasis is “fluid”: higher-order interactions dictate that the epistatic interaction between two mutations depends strongly on the genetic background. Only a small subset of mutations, primarily at functionally critical sites, display signatures of strong global epistasis, while the majority do not. This dichotomy reveals a strikingly “binary” nature of epistasis. Finally, we introduce a novel framework to predict the distribution of fitness effects (DFE) of genotypes: the phenotypic DFE, which offers a statistical route to predicting evolutionary outcomes even when individual mutational effects are unpredictable.

## Results

### *folA* fitness landscape in *E. coli*

Papkou and colleagues recently generated a fitness landscape of a 9-bp region of *folA*, a gene known to be important for resistance to the antibiotic trimethoprim^43,50,54^. They systematically constructed all 4^9^ possible variants in this region, grew them in media containing trimethoprim, and quantified relative fitness using deep sequencing. This approach yielded reliable fitness estimates for ∼99.7% of the variants. Their analysis revealed a highly rugged landscape with 514 fitness peaks. Despite this ruggedness, most genotypes retained adaptive access to high-fitness peaks, largely because such peaks were surrounded by broad basins of attraction.

Using this dataset, we first examined how the structure of the landscape changes with scale. As landscape size increases, the absolute number of peaks rises, but the density of peaks declines (**Supplement Note I; Supplement Figures S2–S3**). At the same time, accessibility of the global fitness peak decreases (**Supplement Figure S4**). Only limited regions of the landscape can be represented as maximally rugged NK landscapes (**Supplement Figure S5**). This pattern highlights a fundamental scaling property of protein fitness landscapes: as dimensionality increases, ruggedness intensifies, making global optima harder to reach. Such constraints are critical for understanding both the limits of protein evolvability and the predictability of adaptive trajectories.

Following Papkou *et al.*, we classify genotypes into two categories: functional (∼7% of all variants) and non-functional (∼93%). This classification was based on statistical segregation of the 4^9^ measured points, and we adopt the same distinction here.

A key limitation of the original analysis is that it relied on a single experimental replicate, without accounting for measurement error. To address this, we re-analyzed the landscape using average fitness values across replicates together with their standard deviations. Incorporating experimental uncertainty substantially alters the inferred structure of the landscape. Of the 514 peaks reported by Papkou *et al.*, only 127 remain when error is considered. Strikingly, all surviving peaks correspond to high-fitness genotypes. In contrast, the non-functional portion of the landscape (>90% of genotypes) contains no peaks at all.

Thus, to minimize bias from any single experimental run, all subsequent analyses were performed using replicate-averaged fitness values together with their associated errors.

### Higher-order epistasis makes interactions between two mutations “fluid”

Epistasis between two mutations can take different forms: no epistasis, positive, negative, or sign epistasis^55^. In the absence of epistasis, the combined effect of two mutations equals the sum of their independent fitness effects. When the combined effect deviates significantly from this sum but retains the same overall direction (beneficial or deleterious), the interaction is classified as positive or negative epistasis. In contrast, sign epistasis occurs when effect of either mutation has the opposite sign from what it exhibits in another genotype. Sign epistasis can be further subdivided into single, reciprocal, and other categories, depending on the number of evolutionary paths that are blocked to Darwinian adaptation.

While individual examples of each type of epistasis are well documented^37,55^, it remains unclear how often the same pair of mutations switches between categories depending on the genetic background. Put differently, is the “nature” of pairwise epistasis itself stable, or does it vary across the landscape?

To address this, we quantified, across all possible genetic backgrounds, the fraction of genomes in which all pairs of mutations exhibit (a) positive epistasis (PE), (b) negative epistasis (NE), (c) sign epistasis [single (SSE), reciprocal (RSE), or other (OSE)], or (d) no epistasis (**Figure 1A**).

**Figure 1.**
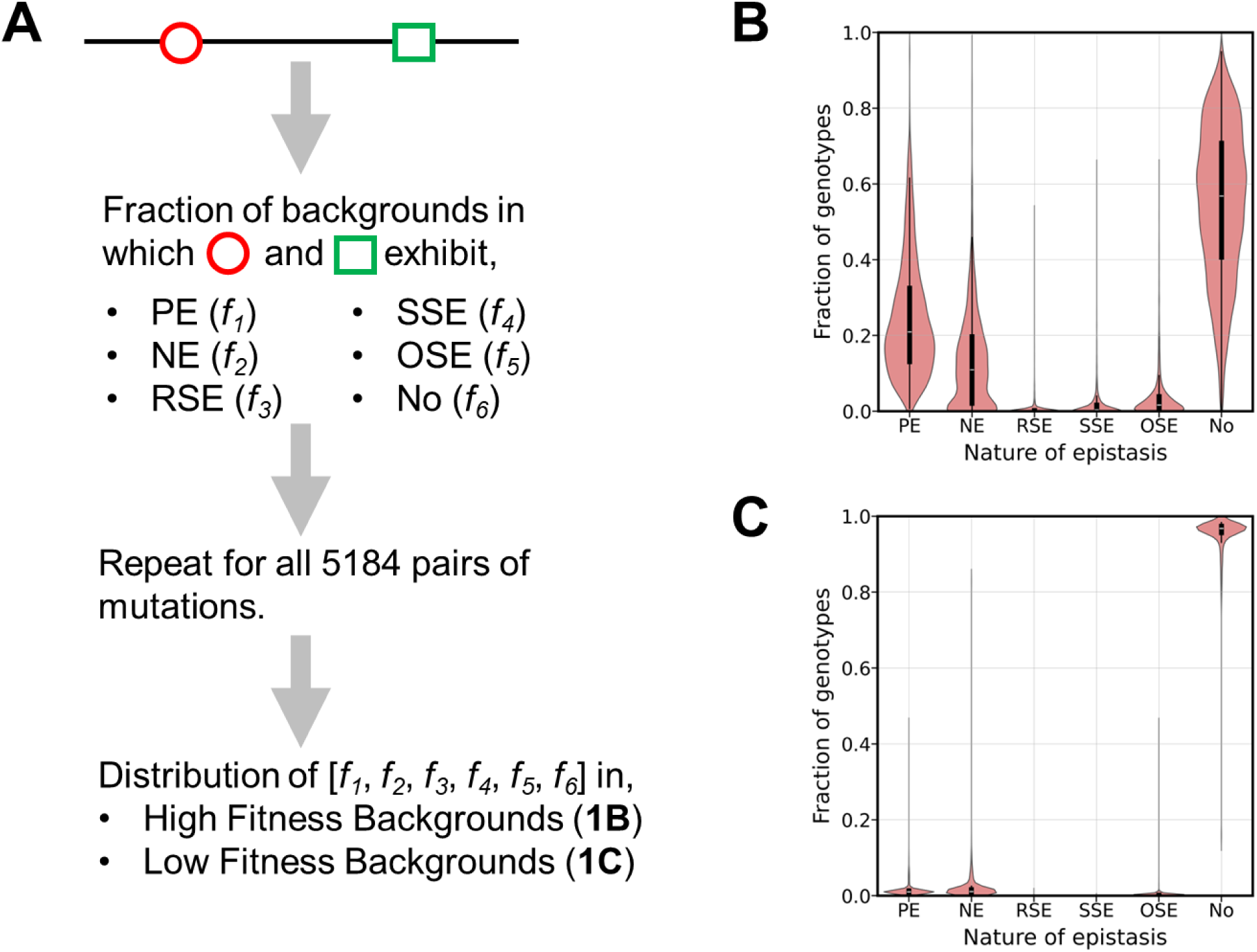
The nature of epistasis between two mutations depends on genetic background. **(A)** Example illustrating how the nature of epistasis between two mutations (red circle and green square) was quantified across all 47 possible genetic backgrounds. For each pair, we calculated the fraction of genomes in which the interaction was classified as positive epistasis (PE), negative epistasis (NE), reciprocal sign epistasis (RSE), single sign epistasis (SSE), other sign epistasis (OSE), or no epistasis. **(B–C)** Distributions of these fractions across all possible mutation pairs in **(B)** functional and **(C)** non-functional backgrounds. Interactions are far more diverse and dynamic in functional backgrounds, whereas in non-functional backgrounds most pairs exhibit no epistasis.

As an example, the mutation pair G→A at position 3 and T→C at position 7 displays striking variability. In functional backgrounds, this pair shows positive epistasis in 12.7% of cases, negative in 9.1%, single-sign epistasis in 0.9%, other-sign epistasis in 1.8%, and no epistasis in 75.2%. By contrast, in non-functional backgrounds, the same pair does not exhibit epistasis in 97.0% cases, with only rare positive or negative epistasis. For ∼0.5% of mutants, fitness data were unavailable. Comprehensive results for all mutation pairs are provided in **Supplement Data File 1**.

We performed this analysis with all 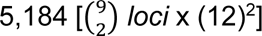 possible mutation pairs in the 9-bp *folA* region. The resulting frequency distributions are shown in **Figures 1B–C** for functional and non-functional backgrounds, respectively. On average, in functional backgrounds, mutation pairs exhibited no epistasis in ∼57% of cases, followed by positive epistasis (∼21%), negative epistasis (∼11%), and rare forms of sign epistasis (≤2%). In non-functional backgrounds, the overwhelming majority of interactions (>96%) showed no epistasis, with positive or negative interactions observed only rarely (**Figure 1C; Supplement Table 1; Supplement Data File 2**).

These results demonstrate that the nature of epistasis between two mutations is not fixed but contingent on the surrounding genetic background. This contingency arises from higher-order epistasis^56–59^, which modulates how pairwise interactions manifest. As a result, pairwise epistatic interactions are “fluid,” shifting in character across the landscape. Importantly, this fluidity is far more pronounced among functional variants than among non-functional ones (**Supplement Data File 1)**.

### Functionally important sites cause more frequent switches in epistasis

Previous work has shown that both the secondary structure and spatial location of a residue can influence the type of epistasis it exhibits^60^. Building on this, and on our finding that the nature of epistasis between two mutations varies across genetic backgrounds (**Figure 1**), we next asked whether some positions are more prone than others to reshaping pairwise interactions. In other words, are certain loci especially influential in driving higher-order epistasis?

To quantify this, we measured how often the introduction of a mutation at a given locus, X, alters the type of epistasis between two other mutations (**Figure 2A**). This analysis reveals that epistatic switching is common in functional backgrounds: adding a single mutation at locus X changes the interaction type in more than 35% of cases (**Figure 2B**). By contrast, in non-functional backgrounds the same measure is typically below 10% (**Figure 2C**). Moreover, not all loci contribute equally. Sites 4 and 5, which are known to be critical for *folA* function, exert the greatest influence, driving far more switches in epistasis than the other positions on the landscape (**Supplement Tables 2–3**). Thus, functionally important residues can govern how interactions between other sites are expressed, a property that may strongly shape protein evolution.

**Figure 2.**
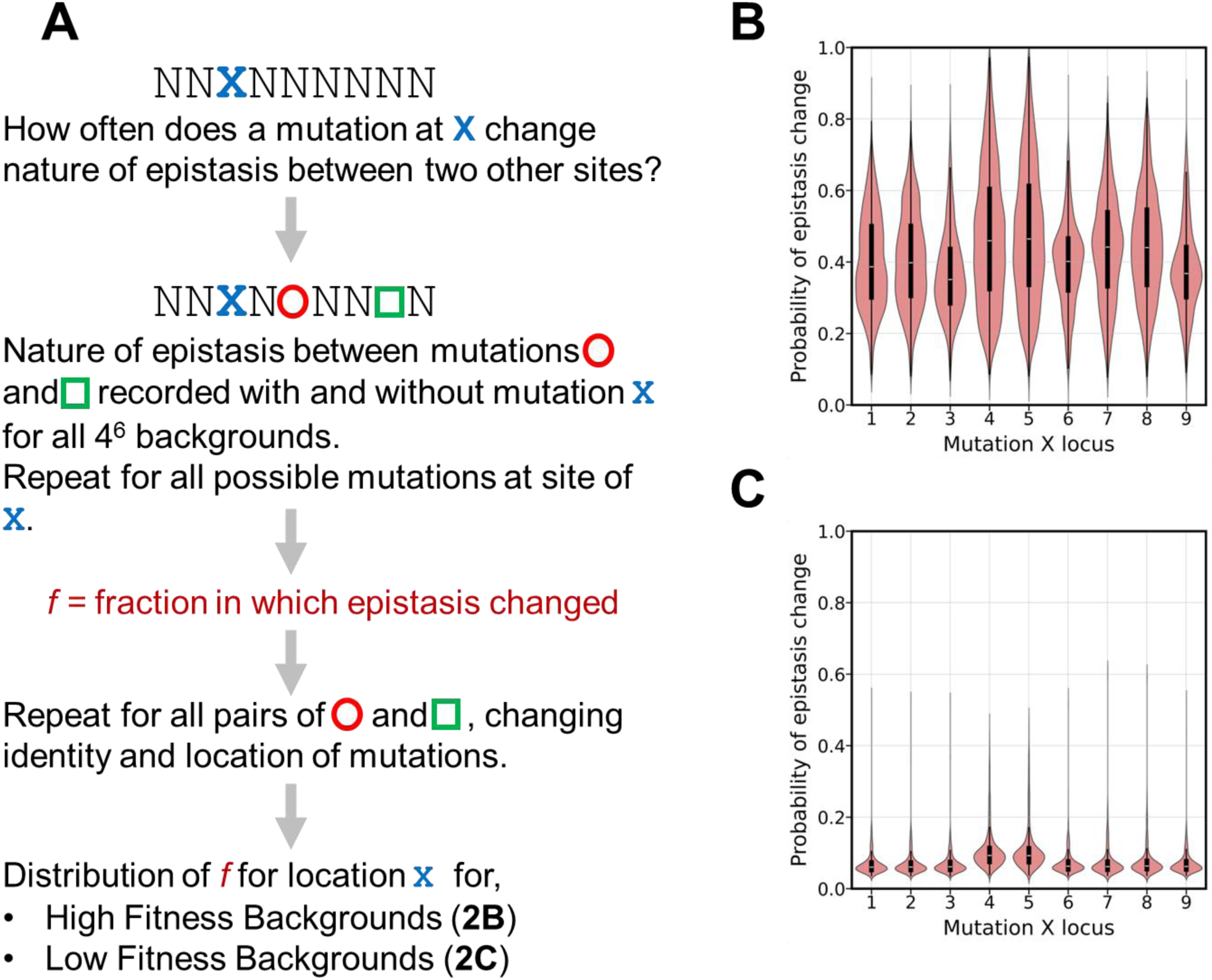
Functionally important sites drive switches in epistasis more often than others. **(A)** Schematic of the approach used to measure the impact of a mutation at locus X on the interaction between two other mutations (red circle and green square). For each pair, the six remaining sites (N) were fixed, and the type of epistasis was recorded before and after introducing a mutation at X. Across all 4^6^ genetic backgrounds, we calculated the fraction *f* of cases in which this mutation altered the nature of epistasis. **(B–C)** Distributions of *f* for each locus in **(B)** functional and **(C)** non-functional backgrounds. Mutations at positions 4 and 5, which are known to be critical for *folA* function, cause switches in epistasis far more frequently than other sites.

The most frequent outcomes of such switches are transitions to no epistasis, positive epistasis, or negative epistasis (**Supplement Figures S6–S7; Supplement Data File 3**). Switches to sign epistasis are less common overall. However, mutations at the functionally important sites (positions 4 and 5) trigger sign epistasis more often than mutations at the other seven positions, both in functional and non-functional backgrounds.

These results highlight two key points. First, higher-order epistasis makes pairwise interactions remarkably fluid, as even a single mutation elsewhere can change their character. Second, this fluidity is not evenly distributed across the genome: functionally important sites disproportionately drive the switching of epistatic interactions. Such site-specific control may complicate the predictability of evolutionary trajectories. Indeed, analysis of a previously published five-point fitness landscape of the β-lactamase gene**^31^** suggests that frequent switching of nature of epistasis is a general feature of intramolecular landscapes (**Supplement Figure S8**).

### Synonymous mutations can alter the nature of epistasis

Although synonymous mutations were long considered neutral^61,62^, many studies now show that they can substantially affect cellular fitness^63–66^. However, their potential role in shaping epistasis, particularly higher-order interactions within a protein, has not been investigated.

To address this question using the *folA* landscape, we asked whether introducing a synonymous mutation at a codon unaffected by a given pair of mutations could change the type of interaction between them (**Figure 3A**). Depending on the position of the pair, one or two codons remain unaltered, and we systematically introduced all possible synonymous substitutions at these codons. For stop codons, we treated TAA TGA and TAA TAG as synonymous (see Discussion).

**Figure 3.**
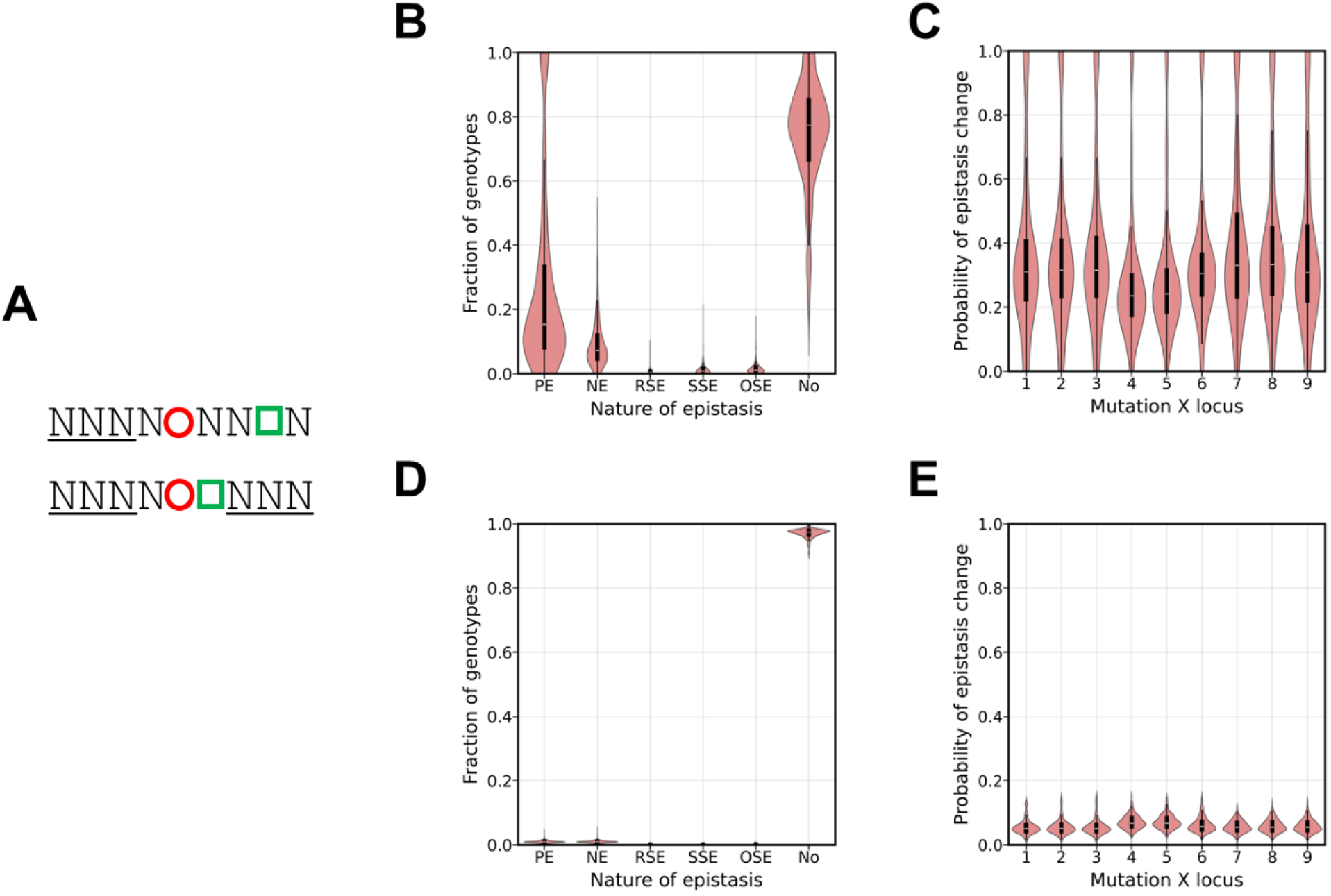
Synonymous mutations can alter the nature of epistasis between two mutations. **(A)** Schematic of the approach: for each pair of mutations (red circle and green square), the codons not directly affected (underlined) were identified, and all possible synonymous substitutions were introduced at these sites. We then assessed whether the type of epistasis between the focal pair changed. **(B–C)** Distributions of epistasis types and switching frequencies in functional backgrounds. **(D–E)** Corresponding distributions in non- functional backgrounds. In functional backgrounds, synonymous substitutions frequently altered the nature of epistasis, although less often than non-synonymous substitutions. In non-functional backgrounds, such changes were rare.

This analysis reveals that synonymous mutations can indeed alter the nature of epistasis, and do so frequently (**Figures 3B–E**). In functional backgrounds, the likelihood of such changes is substantially higher than in non-functional backgrounds. Quantitatively, the frequency of change due to synonymous mutations is statistically lower than for non-synonymous mutations, yet the overall trends are qualitatively similar to those described in **Figures 1–2** (**Supplement Figures S9-S10**).

These findings demonstrate that even synonymous substitutions can reshape pairwise interactions, further reinforcing the “fluid” character of epistasis. This pervasive fluidity complicates prediction of evolutionary trajectories. Nonetheless, epistasis is known to follow certain statistical regularities, collectively termed “global” patterns^1^. We therefore next examined whether these global patterns are also evident in the *folA* landscape.

### Non-synonymous mutations, but not synonymous mutations, on the landscape exhibit diminishing returns and increasing costs

While synonymous substitutions highlight the fluid and context-dependent nature of local interactions, a different picture can emerge at larger scales. Specifically, many protein landscapes exhibit broad statistical regularities in how mutations contribute to fitness, such as diminishing returns^1,67–69^. We therefore next examined whether such global patterns are also evident in the *folA* landscape. One such pattern is global epistasis, where the effect of a mutation systematically depends on the fitness of the background genotype. Typically, beneficial effects weaken as background fitness rises (diminishing returns), while deleterious effects become more severe in higher-fitness genotypes (increasing costs) (**Figure 4A**)^10,70,71^.

**Figure 4.**
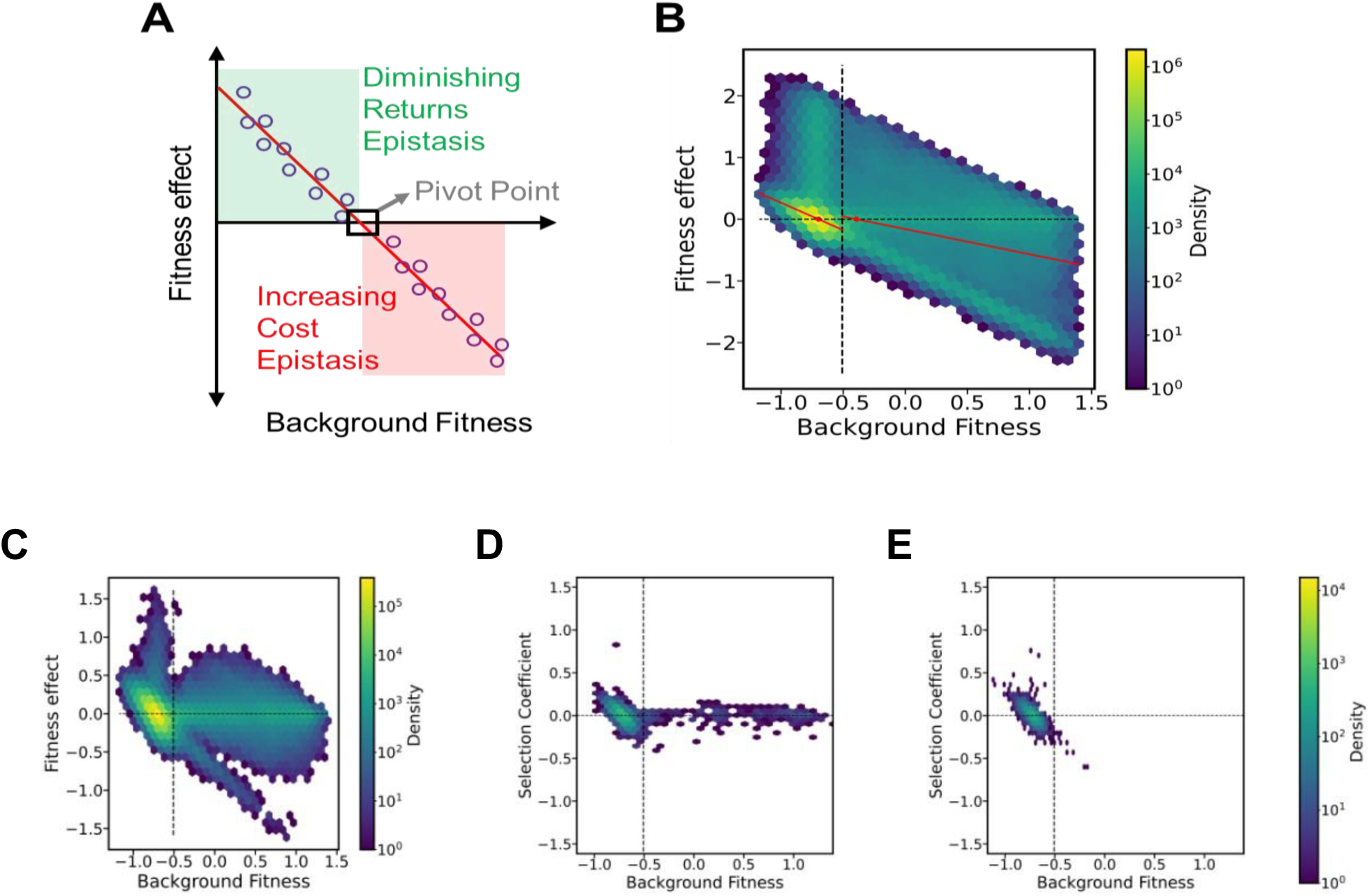
The folA fitness landscape exhibits weak patterns of global epistasis, which are absent for synonymous mutations. **(A)** Cartoon illustrating global epistasis: the beneficial effect of a mutation decreases as background fitness increases (diminishing returns epistasis). Beyond a pivot point, the same mutation becomes deleterious, and its negative effects increase with higher background fitness (increasing costs epistasis). **(B)** Empirical test for global epistasis: mutational effects (y-axis) plotted against background fitness (x-axis). A weak but significant negative trend is observed in both low-fitness (R² = 0.1236; left of dotted line) and high-fitness (R² = 0.1284; right of dotted line) backgrounds. Red lines show best linear fits. **(C)** Synonymous mutations show no such correlation between effect size and background fitness. **(D–E)** Instead, most synonymous mutations fall into two recurrent patterns. Examples are shown for the synonymous substitution AAA → AAG at position 1 **(D)** and at position 2 **(E)**. Full plots for all synonymous substitutions are provided in the Supplement.

On the *folA* landscape, plotting mutational effects against background fitness reveals weak signatures of global epistasis (**Figure 4B**). In both low- and high-fitness regions of the landscape, the negative slope of the relationship is consistent with diminishing returns and increasing costs. However, the correlations are modest (R² = 0.1236 for non-functional backgrounds; R² = 0.1284 for functional backgrounds), indicating that, overall, global epistasis is not very strong on the *folA* landscape. However, there is a clear distinction in the slopes of all mutations. The lethal mutations have a slope smaller than -0.7 and average slope of -0.98. The remaining mutations all have a slope greater than -0.56.

Synonymous mutations, however, break this pattern. When their effects are plotted against background fitness, no correlation is observed (R^2^ < 0.1) (**Figure 4C**). Instead, synonymous substitutions tend to follow two recurring behaviors, which appear across many codons. In the first, the effect of a mutation is independent of the background fitness **(Figure 4D)**, and in the second, non-functional backgrounds show a negative correlation between fitness effect and background fitness **(Figure 4E)**. (see Supplement for full set). Thus, unlike non-synonymous mutations, synonymous changes do not exhibit very strong background fitness-dependence. Alternatively, the fitness effect readouts of synonymous mutations are small and therefore, detection of signatures of global epistasis difficult.

There are, nevertheless, a few notable exceptions. GAN at locus 4 generates a fraction of non-functional sequences; stop codon conversions such as TAG → TAA at codons 1 or 3 are lethal, while the reverse TAA → TAG restores function. Additionally, synonymous substitutions at codons TCA or TCG in codons 1 or 3 yield unusually high selection coefficients in high-fitness backgrounds. These outliers highlight the diversity of synonymous effects but reinforce the broader conclusion: while non-synonymous mutations conform (weakly) to global epistasis, synonymous mutations escape it.

We next tested whether the average fitness effect of a synonymous mutation correlates with the frequency with which it alters the type of epistasis between two mutations. No such relationship was observed (R² ≈ 0.11; **Supplementary Figure S11**), indicating that the ability of synonymous mutations to reshape epistasis is not simply a byproduct of their direct fitness effects.

### Only a small fraction of mutations exhibit strong global epistasis

We next examined the fitness effects of all 108 possible mutations (12 per site × 9 sites) across the *folA* fitness landscape. As shown in **Figure 5**, only a small subset of mutations exhibits strong global epistasis (R^2^ > 0.5)^72^. 14 of the 16 such cases occur at nucleotides 4 and 5, which correspond to catalytically important positions *folA*^46^. This indicates that the drivers of global epistasis are not distributed uniformly across the gene but instead concentrated at functionally critical sites. The majority of mutations (77/108) do not show a strong correlation (R² < 0.2) between their fitness effect and the background fitness (**Supplement Figure S12, Supplement Table S4**). Here we focus only on whether such a correlation with background fitness exists, rather than on the slope or magnitude of the relationship. Our goal is to capture the presence or absence of strong global patterns in a generic, statistical sense, rather than to quantify the precise strength of each interaction.

**Figure 5.**
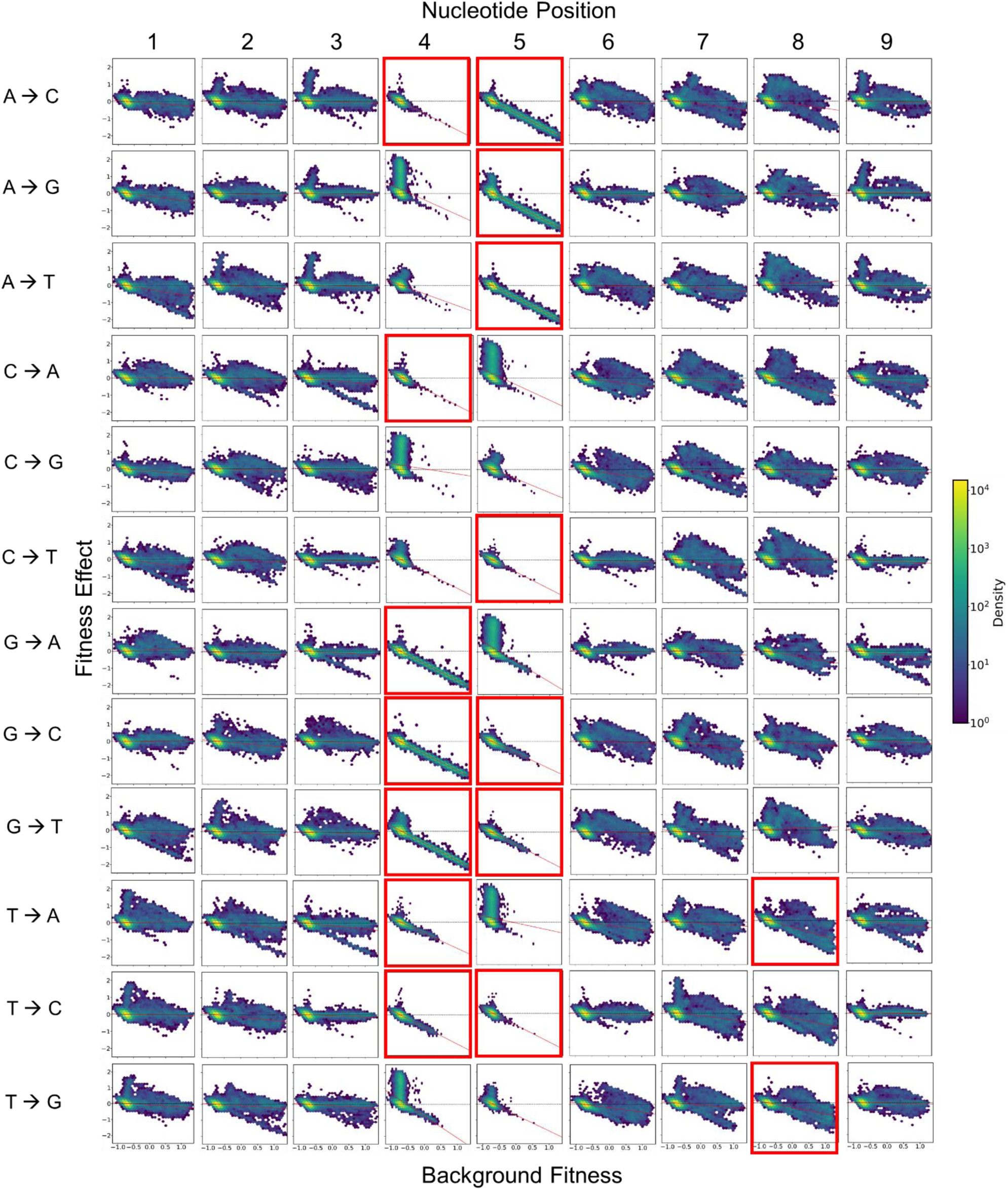
Only a small fraction of mutations exhibit strong global epistasis. Among the 108 possible mutations in the *folA* landscape, only 16 exhibit statistically significant patterns of global epistasis (R² > 0.4). Strikingly, 14 of these are located at nucleotide positions 4 and 5 (highlighted in red boxes), which are functionally critical for *folA*. Complete statistics for all mutations are provided in Supplementary Table S4.

This sharp contrast suggests a binary behavior of mutations: they either exhibit strong global epistasis (R^2^ > 0.5), or not (R^2^ < 0.5). In *folA*, strong global epistasis is essentially restricted to mutations at functionally crucial positions. Such “binary” global epistasis (where mutations are predicted to exhibit strong global epistasis at a few sites and only weakly exhibit global epistasis at other sites) has recently been predicted in a theoretical model of intermolecular epistatic patterns^73,74^. Our results suggest that such patterns exist in intramolecular epistasis too.

Beyond this binary classification, a recent study^75^ showed that pivot growth rate, or the fitness at which the sign of the fitness effect of a mutation switches, is conserved across mutations for an environment, implying an underlying cellular constraint. We tested this on the *folA* landscape and found that ∼80% of mutations pivot around a growth rate of −0.657 ± 0.0657 (**Figure 6, Supplement Table S4**). Importantly, the pivot emerges even for mutations that otherwise show weak global epistasis. Thus, the pivot growth rate represents a conserved organizing principle of mutational effects, beyond the globality of epistasis.

**Figure 6.**
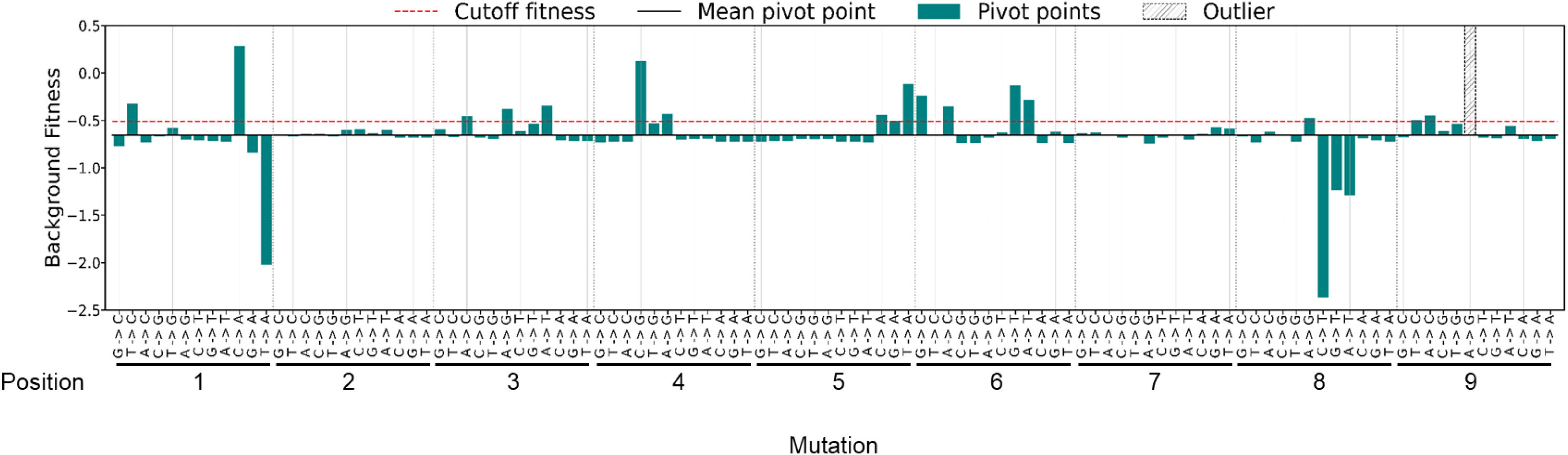
Most mutations pivot from beneficial to deleterious effects at a common fitness threshold. Background fitness (y-axis) at which each of the 108 mutations (x-axis) switches from being beneficial to deleterious is shown. The solid black line marks the average pivot fitness (-0.657), while individual bars represent the deviation of each mutation from this mean. More than 80% of mutations (86 out of 108) pivot within the range -0.657 ± 0.0657. The red dotted line indicates the growth rate used in Papkou et al. ^40^ to distinguish functional and non-functional variants.

### Predicting the distribution of fitness effects (DFE) from phenotype

The presence of a shared pivot illustrates that mutational effects can sometimes be captured by simple, robust principles. However, such principles appear to be the exception rather than the rule. As shown earlier, only mutations at functionally important sites exhibit highly global behavior, while the majority display idiosyncratic, background-dependent effects. Consequently, even though pivots highlight one predictable axis of mutational behavior, the overall distribution of fitness effects (DFE) remains difficult to predict from phenotype alone. Without prior knowledge of functionally constrained sites or the specific genetic background, forecasting the outcomes of mutations across the landscape remains unreliable.

We therefore asked whether any higher-level statistical patterns might nevertheless enable prediction. To this end, we defined the phenotypic DFE, the average distribution of fitness effects across all genotypes of similar fitness (**Figure 7A**). To compute this, we binned genotypes into narrow, non-overlapping fitness intervals of width 0.05. For each genotype, the DFE was obtained by introducing all 27 possible point mutations (three alternative nucleotides at each of the nine sites). These DFEs were then averaged within each interval to obtain the phenotypic DFE. We next tested whether the phenotypic DFE could reliably predict the DFE of individual genotypes of comparable fitness.

**Figure 7.**
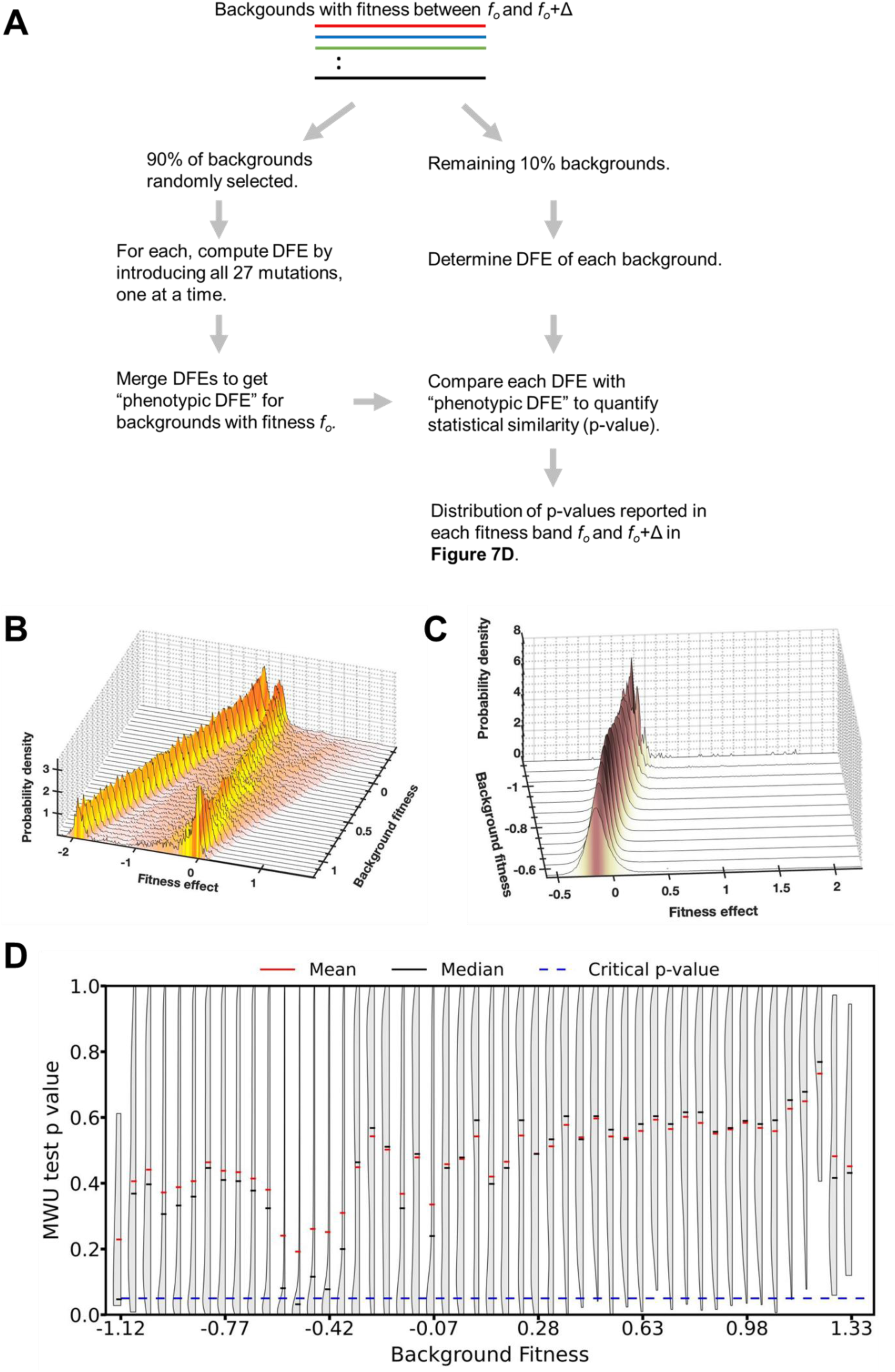
Phenotypic DFE captures robust, fitness-dependent patterns. **(A)** To define a “phenotypic DFE,” genotypes of near-identical fitness were split into two groups (90% and 10%). The mean DFE of the 90% group was compared with the DFEs of individual genotypes in the 10% group using the Mann–Whitney U test, generating a distribution of p-values. This procedure was repeated across fitness bins. Δ in this work was chosen to be 0.05. **(B)** Phenotypic DFEs of high-fitness backgrounds were bimodal: one peak centered near neutrality (∼0) and a second comprising deleterious mutations whose magnitude increased with background fitness. **(C)** Phenotypic DFEs of low-fitness backgrounds were unimodal, with the mean shifting towards more deleterious values as fitness decreased. **(D)** Fraction of genotypes in each fitness bin whose DFEs differed significantly (p < 0.05) from the phenotypic DFE. As fitness increases, the phenotypic DFE becomes a progressively better predictor of the true DFE.

To assess predictive power, we computed the phenotypic DFE from 90% of genotypes in a fitness bin, then compared it against the DFEs of the remaining 10%. Similarity was quantified using the Mann–Whitney U test, generating a distribution of p-values for each fitness interval.

The phenotypic DFE exhibited distinct shapes depending on whether backgrounds were functional or non-functional. For functional backgrounds, the DFE was bimodal: a strong peak of large deleterious mutations whose effect intensified with increasing background fitness, and a second peak centred near neutrality (**Figure 7B**). For non-functional backgrounds, the DFE was unimodal, with the mean shifting towards increasingly deleterious values as background fitness increased (**Figure 7C**).

Crucially, the phenotypic DFE was more predictable from background fitness than the effects of individual mutations. As background fitness increased, the fraction of genotypes whose DFEs were statistically distinguishable from the phenotypic DFE decreased (**Figure 7D**). This trend is also visible in the distribution of p-values across fitness bins (**Supplement Figure S13**). Thus, while individual mutational effects are unpredictable, higher-level patterns captured by the phenotypic DFE emerge as statistically robust and fitness-dependent.

## Discussion

Our systematic mapping of the *folA* fitness landscape highlights how epistasis is both pervasive and fluid, reshaping the way mutations contribute to fitness depending on the genetic background. While previous work has shown that strongly deleterious mutations can interact in unpredictable ways^76,77^, our results demonstrate that even synonymous and stop-codon variants, which are often assumed to be functionally neutral or uniformly disruptive, can substantially alter the architecture of epistasis. This underscores the importance of considering all classes of mutations when evaluating the genotype–phenotype–fitness map.

At the molecular level, the mechanisms behind these interactions are multifaceted^78–83^. Mutations at catalytic residues predictably exert large effects on enzyme activity, but their fitness impact is strongly modulated by the presence of other substitutions^84–87^. For instance, alleles that increase catalytic efficiency exhibit diminishing returns epistasis when combined with overexpression mutations: once enzyme activity is already high, further increases yield smaller fitness gains^81,88,89^. This scaling behavior illustrates how basic biochemical principles, such as saturation of substrate conversion, give rise to global patterns of epistasis.

By contrast, synonymous mutations and stop codons likely most frequently act through RNA-level features^90–93^. Synonymous substitutions can alter codon usage, mRNA folding, or translation rates, thereby modifying protein abundance^90,94^. These changes shift the effective catalytic capacity of the enzyme, which in turn modulates the fitness consequences of subsequent coding changes^49,95,96^. Similarly, premature stop codons, while truncating the protein, may still influence transcriptional or translational dynamics in ways that reshape the adaptive potential of other mutations^97–101^. Such findings expand the traditional view of epistasis beyond amino acid–level interactions, highlighting the integrated contributions of DNA, RNA, and protein properties to fitness.

Importantly, we find that epistasis at *folA* is not purely local but also exhibits global regularities. The systematic shift in the distribution of fitness effects with background fitness reflects a general principle: as genotypes approach higher catalytic efficiency and thus higher fitness, the availability of beneficial mutations decreases. This mirrors patterns observed in long-term microbial evolution experiments, where initial rapid gains are followed by slower adaptation^102,103^, and suggests that our short-term, locus-specific landscape captures dynamics relevant at the scale of whole genomes and populations.

Overall, these results emphasize two key insights. First, the context dependence of mutations means that adaptive outcomes cannot be predicted by examining mutations in isolation: the same change may be beneficial, neutral, or deleterious depending on background^104,105^. Second, the scaling behavior of epistasis reveals that despite this contingency, evolutionary dynamics are governed by reproducible rules rooted in molecular biophysics^106,107^. By tying catalytic efficiency, protein abundance, and RNA-level features to fitness outcomes, our work shows how molecular details carve the shape of adaptive landscapes^63,80^. This integrated perspective not only clarifies the mechanistic basis of epistasis at *folA* but also provides a template for understanding how genetic, regulatory, and structural layers interact to shape evolutionary trajectories.

## Methods

### Calculating Hamming Distance

Hamming distance between two sequences of equal length was calculated by comparing the sequences and counting the number of loci with differing nucleotides.

### Construction of Sequence Spaces and Landscapes

To construct an *n*-base pair sequence space from a parent *p*-base pair sequence space (*p* > *n*), we fixed *p* − *n* loci in the parent sequence space and enumerated with the selected loci held constant. By iterating over all 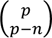 combinations of fixed loci and all 4^*p*−*n*^ permutations of their sequence, the parent sequence was partitioned into 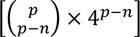 *n*-base pair subspaces (**Supplement Table 2**).

For this study, a nine-base pair parent sequence space was used to generate sequence spaces with 1 ≤ *n* ≤ 9. Fitness landscapes were constructed by mapping the empirical fitness values of sequences in each subspace to the corresponding sequences in the Papkou et al DHFR dataset^43^. Variants lacking fitness data were excluded from the analysis.

### Identifying Peaks in a Fitness Landscape

#### Number of Peaks

Peaks were defined as variants with fitness higher than all their one-Hamming-distance neighbors.

#### Peak Probability

The ratio of peaks to total sequences in a landscape.

#### Expected number of peaks

For a four-letter genome, expected peaks in a maximally rugged NK landscapes is given by 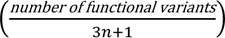 for an *n*-base pair landscape^20^. Results were rounded off to the nearest integer.

### Fitness Effects of Mutations

#### Mutational Effects

For a mutation acting on a genotype, the fitness effect *s* = *f*_*m*_ − *f*_*b*_, was calculated where *f*_b_ is the background fitness and *f* is the mutant fitness.

#### Genotypic DFE

For any 9-bp sequence, all 27 possible single-base mutations were evaluated to construct the genotype-specific DFE.

#### Phenotypic DFE

DFEs of all genotypes within a narrow fitness interval (width 0.05) were averaged to generate a phenotypic DFE.

#### Statistical Tests

Mann–Whitney U test and Kolmogorov–Smirnov tests were performed using using the *python scipy.stats* library.

#### Linear Regression

Pivot points for mutations were estimated by linear regression of background fitness versus selection coefficient using the “LinearRegression” model from python sklearn.linear_model library.

### Epistasis as Function of Genetic Background

#### Classifying Epistasis

For two mutations *a* → *A* and *b* → *B* fitness effects *S*_*A*_, *S*_*B*_, and combined effect *S*_*AB*_ were computed. Variance was calculated by assuming independence of fitness measurements, with six replicates per sequence.

#### Hypothesis testing

Four hierarchical t-tests (p = 0.05) classified mutation pairs into:

- No epistasis: *S*_*AB*_ – (*S*_*A*_ + *S*_*B*_) = 0
- Magnitude epistasis: *S*_*AB*_ = 0
- If *S*_*A*_ = *S*_*B*_, then mutations cannot be classified as exhibiting single sign epistasis
- If *S*_*A*_ = 0, then in combination with the above point, the mutations cannot be classified as Reciprocal Sign Epistasis

#### Epistasis Change with Background

For each mutation pair, genetic backgrounds were compared to their one-mutant neighbors at loci unrelated to the pair (21 neighbors per background) to quantify changes in epistasis.

#### Epistasis Change with Genetic Background

Having the epistasis dossier generated for all mutation pairs, we compiled all cases of *Positive, Negative, Sign* and *No Epistasis*. For each of these cases, we select the genetic backgrounds and their one mutant neighbours such that their differing mutation locus is unrelated to the loci involved in Epistasis (In our case, we can find (9 – 2)*loci* × 3*bases* = 21 such neighbours for each background).

### Sequence Space Traversal

Shortest paths between sequences were enumerated by:

- Listing all mutations required to convert starting sequence *A* to *T*.
- At each step, *s*, collecting sequences at Hamming distance *s* from *A* and ℎ − *s* from *T*, where ℎ is the minimum number of mutations.
- Recursively computing all ℎ! permutations allowing only single-base changes per step.

### Synonymous Mutations

#### Identification

Codons encoding the same amino acid or stop function were enumerated. Synonymous mutations were those differing by a single nucleotide (Hamming distance = 1).

#### Effect on Epistasis

For each mutation pair, unaltered codons were identified, and all synonymous mutations were introduced. The number of instances where the nature of epistasis changed was quantified across all backgrounds and separately for high- and low-fitness subsets.

### Codes

All codes used in this work and Supplement Data Files are available at: 10.5281/zenodo.17012893. The “readme” file at the repository gives details of how to run codes.

## Acknowledgements.

We thank Christian Landry and Krishna Swamy for feedback on the manuscript.

