## Supplementary Figures for "Fluidity and Predictability of Epistasis on an Intragenic Fitness Landscape"

#### Supplement Figure S1.

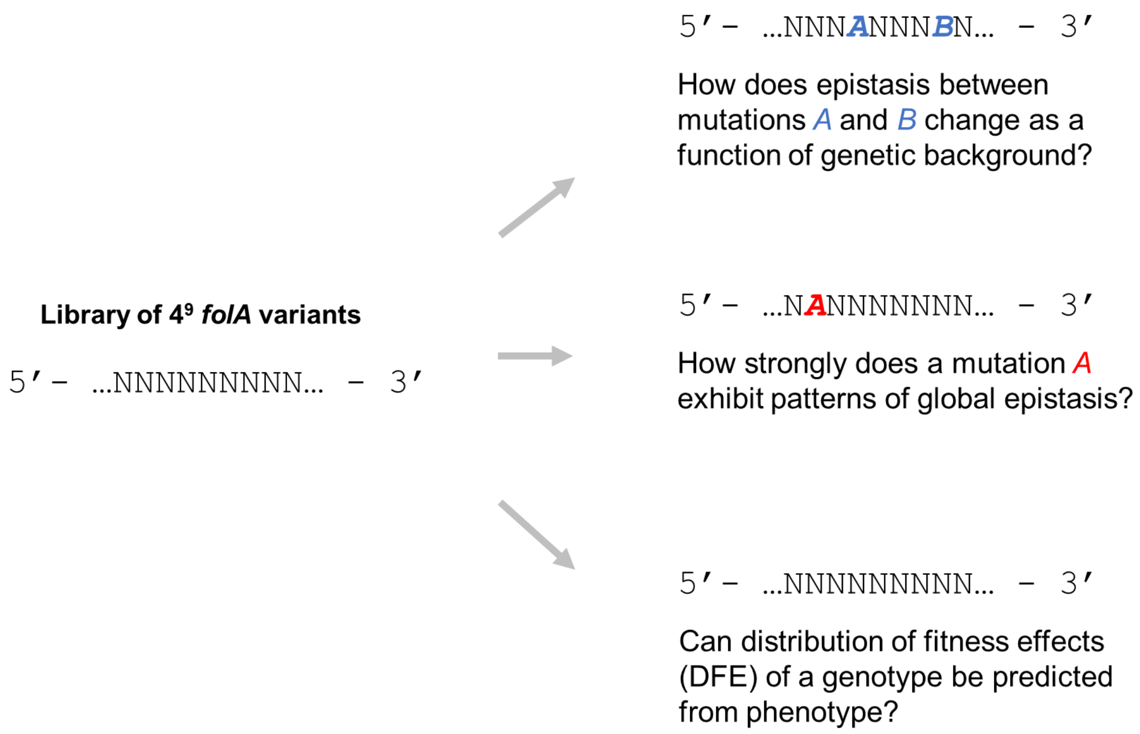


**Figure S1.** Using the data in Papkou and coworkers[1], we use the *folA* landscape to ask three questions. **(A)** How does the nature of epistasis between two mutations (indicated as *A* and *B* in the Figure) change as a function of the genetic background? **(B)** Individually, do all mutations exhibit patterns of global epistasis? **(C)** In the face of idiosyncratic nature of epistasis, can we predict distribution of fitness effect (DFE) of a genotype from its phenotypic traits?

#### Supplement Note I. Analysis of the *folA* landscape.

The number of peaks increases and the probability of a peak decreases, as a function of sequence length. We create subgraphs from a 9-base landscape, so subgraphs of the same size differ in numbers of peaks, resulting in a distribution for each *n*-base pair landscape ($\boldsymbol{n} < 9$). The mean, median statistics of the peaks of these landscapes for each value of *n* is shown. Papkou and coworkers group all sequences into two – "functional" and "non-functional".

We use a maximally rugged (uncorrelated) NK-landscape model to predict the expected number of peaks in a *n*-base pair landscape. Our results show that while the median number of peaks in the landscape distribution is a good predictor of the number of peaks, a large number of landscapes, for any size *n*, do not follow the predicted NK peak distributions.

As *n* increases, while the size of the landscape and the number of peaks both increase exponentially, although the increase in the number of peaks is much slower. Hence, the probability of any given point being a peak decreases with increasing *n*. Our results also show that as the size of the landscape increases, the variation between different subgraphs of the same size decreases, and makes structure of landscapes statistically predictable.

The study by Papkou et al computed that 76.5% of low-fitness variants had access to high-fitness peaks, making the rugged landscape navigable. Theoretically, as we move *n* bases away from a peak on the fitness landscape, the number of shortest trajectories to the peak increases factorially (as *n!*). Though 76.5% of the variants reach peaks through random walks, the accessibility of the peak through all shortest paths is not known.

To explore this, we assessed the accessibility of the highest peak on the landscape for all variants at *n* Hamming distance. The fraction of variants which could not access the peak via even a single shortest path increased from 15% for *n* = 4, to 21% for *n* = 7. These results demonstrate that even with increasing dimensionality of the landscape, sign-epistasis poses an increasing constraint for Darwinian movement on a landscape.

In addition, the peak accessibility of variants at 2 hamming distance from the highest fitness peak in the *folA* landscape suggests that while the fraction of variants which exhibit no sign epistasis (magnitude epistasis) remain the same for low and high fitness backgrounds, the fraction of variants exhibiting reciprocal sign epistasis (no accessible paths to the highest peak) is much greater in high fitness backgrounds than in low fitness backgrounds.

#### Supplement Figure S2.

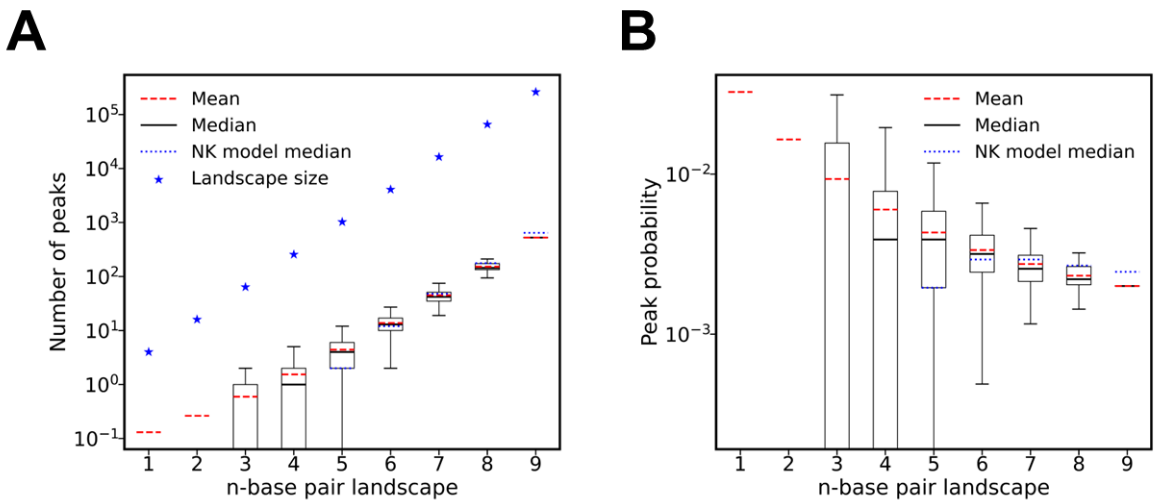


**Figure S2. Probability of peaks in landscapes decreases with size. (A)** The number of high fitness peaks increase exponentially as we increase the size of the landscape, albeit at a slower rate than the increase of size of landscape. **(B)** The probability of a randomly picked point being a high fitness peak decreases with increase in the size of the landscape.

#### Supplement Figure S3.

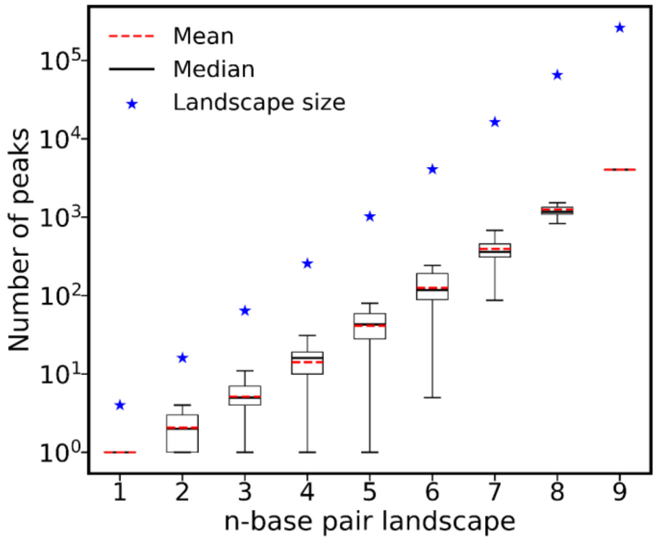


**Figure S3. Total number of peaks in landscape increase at a lower rate than the number of variants in the landscape.** The total number of peaks in the landscape (including high and low fitness peaks) increase exponentially as the landscape size increases, although, much slower than the size of the landscape. Although the functional peaks contribute only a fraction of the total peaks for all size landscapes (median 11.6%; standard deviation 0.938%), this result remains qualitatively same as observed in functional peaks in the main text. Hence, we can conclude that the probability of encountering any fitness peak diminishes as the size of the landscape increases.

#### Supplement Figure S4.

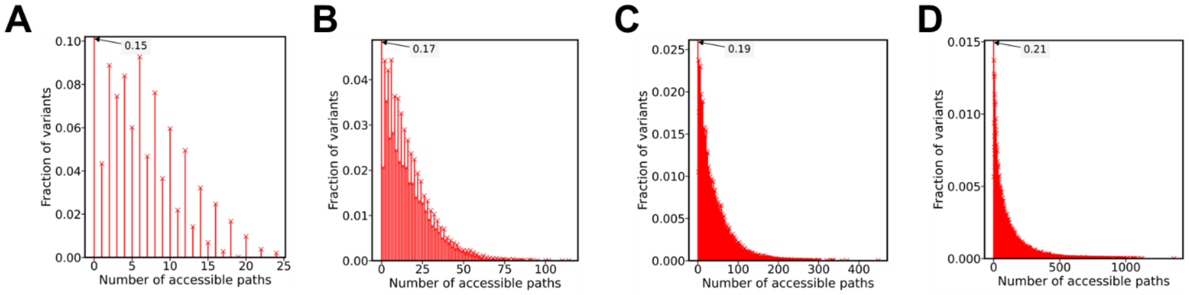


**Figure S4. Path accessibility distribution to highest fitness peak for Hamming distance (A) 4, (B) 5, (C) 6, and (D) 7.** As the Hamming distance increases (A to D), the fraction of points with no adaptive paths to the global peak increases from 0.15 to 0.21.

### Supplement Figure S5.

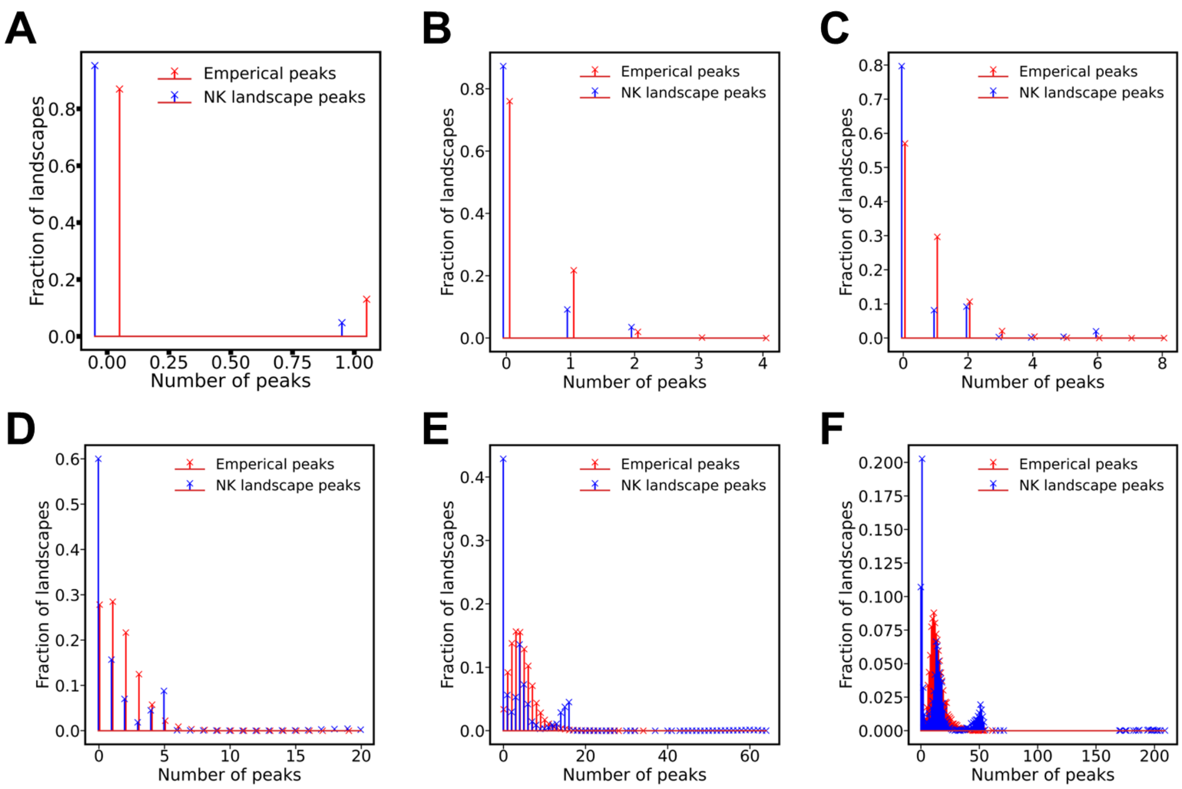


**Figure S5. A large fraction of landscapes does not follow the predicted distribution of maximally rugged NK-model landscape.** Using the number of functional variants present in each landscape, we predicted the expected the number of peaks in maximally rugged (uncorrelated) NK landscape[2]. The expected number of functional peaks in a maximally rugged NK landscape can be found as $\frac{\mathcal{S}}{(A-1)\times L+1}$, where $\mathcal{S}$ is the number of functional variants in the landscape, $A = 4$ is the alphabet size and $L$ denotes the number of sites involved in the landscape. The resulting NK probability distribution (blue stems) is plotted along with the empirical distribution (red stems). Plots A-F are plotted for different landscape sizes: (**A**) 1 nucleotide landscapes; (**B**) 2 nucleotide landscapes; (**C**) 3 nucleotide landscapes; (**D**) 4 nucleotide landscapes; (**E**) 5 nucleotide landscapes; (**F**) 6 nucleotide landscapes. We observed that while the resulting distributions have some resemblance to one another, the p-values of Mann-Whitney U test of the two distributions firmly reject the null hypothesis (**Supplement Table S5**). Hence, contrary to the claim made by Papkou et al[1] (using the single 9 nucleotide landscape), the smaller DHFR fitness landscapes do not resemble the maximally rugged (uncorrelated) NK landscape.

#### Supplement Figure S6.

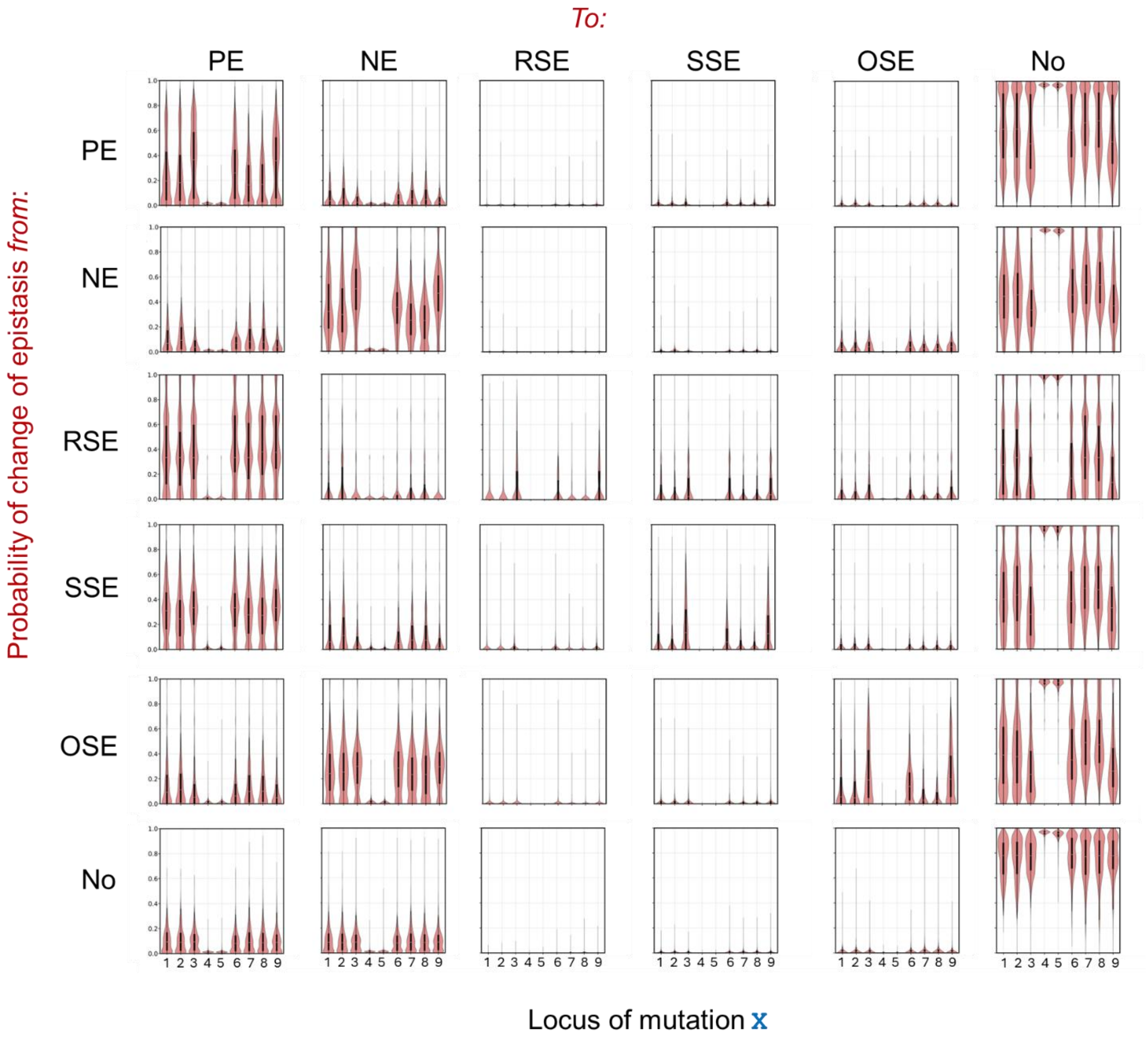


**Figure S6. Change of epistasis as a function of locus *X* in high fitness backgrounds.** Diagonal graphs represent the probability of nature of epistasis not changing. PE: Positive Epistasis; NE: Negative Epistasis; RSE: Reciprocal Sign Epistasis; SSE: Single Sign Epistasis; OSE: Other Sign Epistasis; No: No Epistasis. The probability for change of epistasis was calculated as described in Figure 2A in the main text.

### Supplement Figure S7.

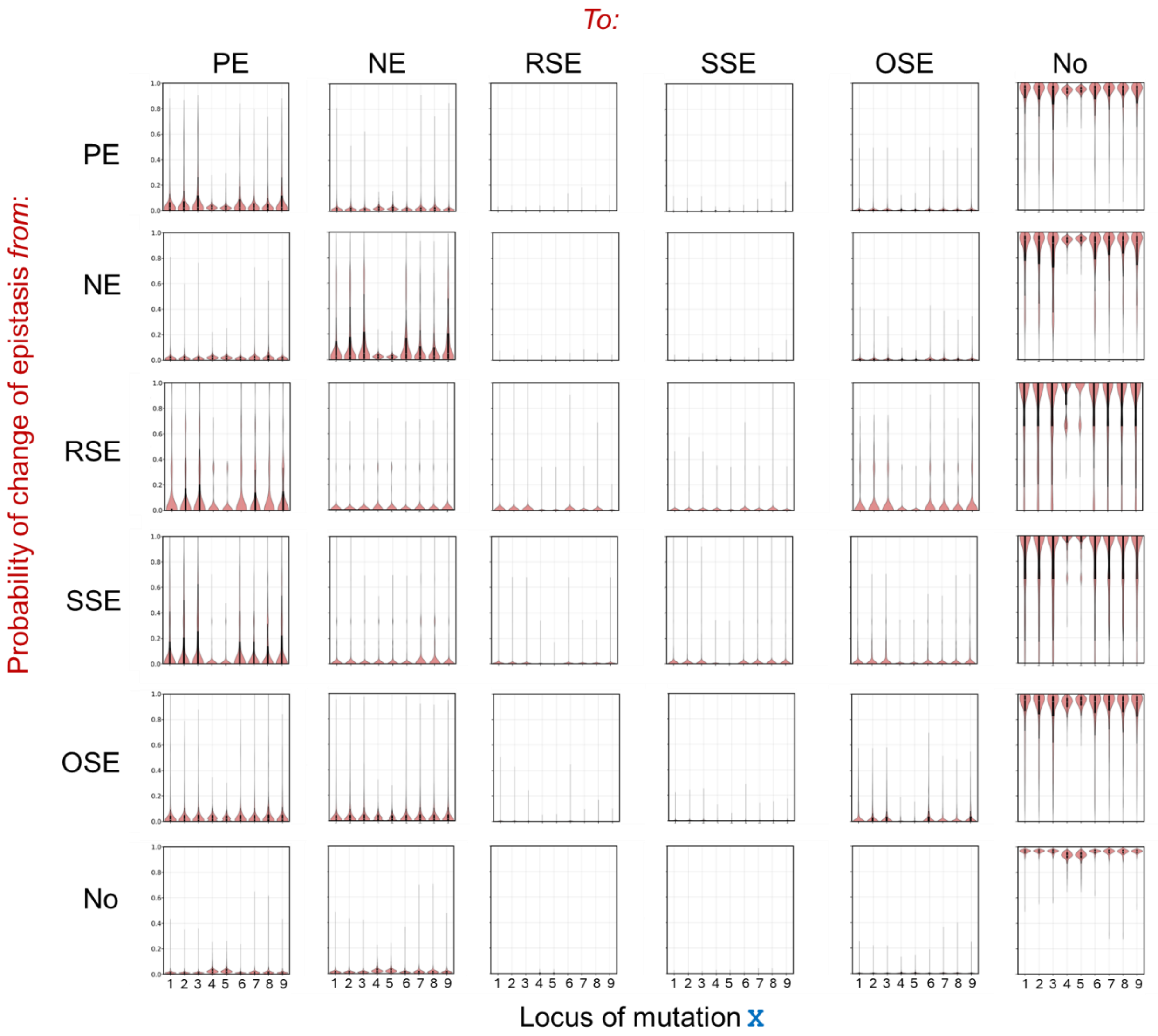


**Figure S7. Change of epistasis as a function of locus *X* in low fitness backgrounds.** Diagonal graphs represent the probability of nature of epistasis not changing. PE: Positive Epistasis; NE: Negative Epistasis; RSE: Reciprocal Sign Epistasis; SSE: Single Sign Epistasis; OSE: Other Sign Epistasis; No: No Epistasis. The probability for change of epistasis was calculated as described in Figure 2A in the main text.

#### Supplement Figure S8.

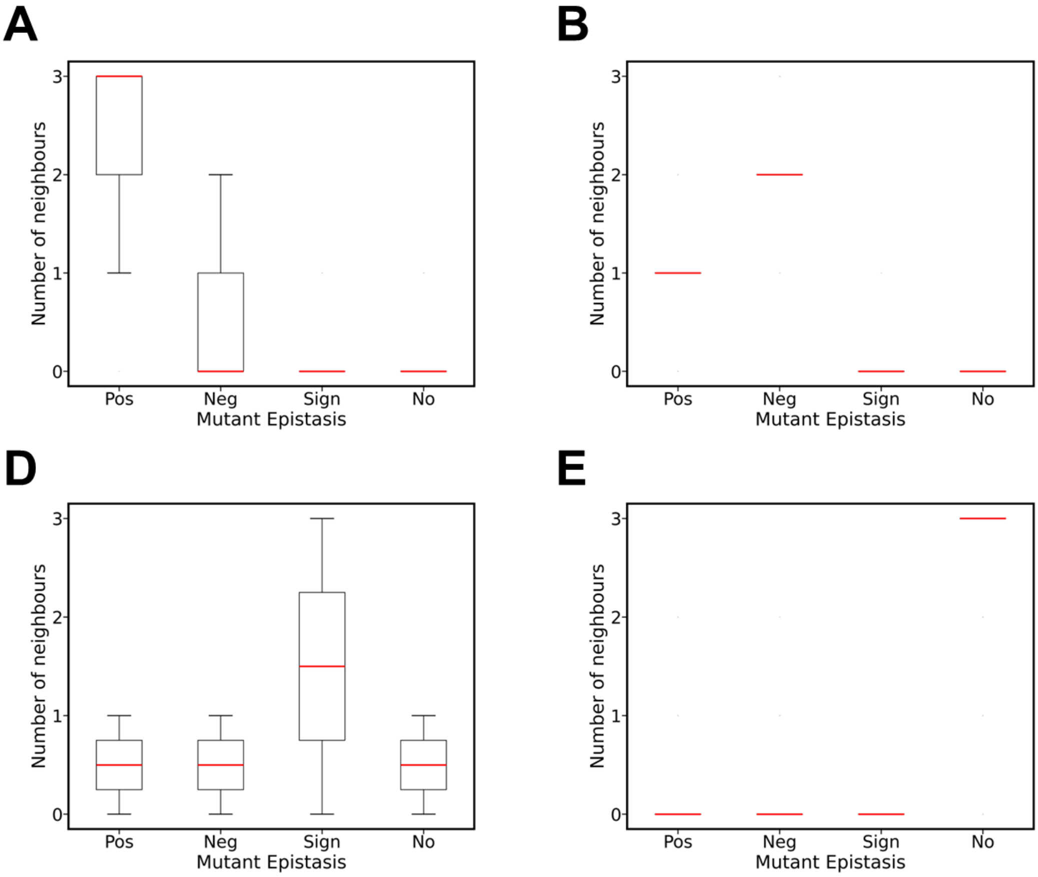


**Figure S8. Epistasis change due to a third mutation observed in beta-lactamase gene.** The phenomenon of nature of epistasis between two loci changing due to a mutation at a third locus is not limited to DHFR gene. In the previously established 5 nucleotide beta-lactamase landscape[3], we noted similar observations. In several backgrounds, the nature of epistasis between two loci were found to change upon a mutation taking place on a third loci. The box plots were created using each mutation pair, for which compiled the cases for variants exhibiting (**A**) Positive epistasis, (**B**) Negative epistasis, (**C**) Sign epistasis and (**D**) No epistasis. The x-axis in the plots represent nature of epistasis acquired by the mutation pairs on the mutant (neighbour), and the y-axis represents the number of neighbours exhibiting the type of epistasis. While positive and negative epistasis occasionally interchange due to a third mutation ans sign epistasis frequently changed to any other form of epistasis due to a mutation, no epistasis changes changes to some other form only in some rare cases.

#### Supplement Figure S9.

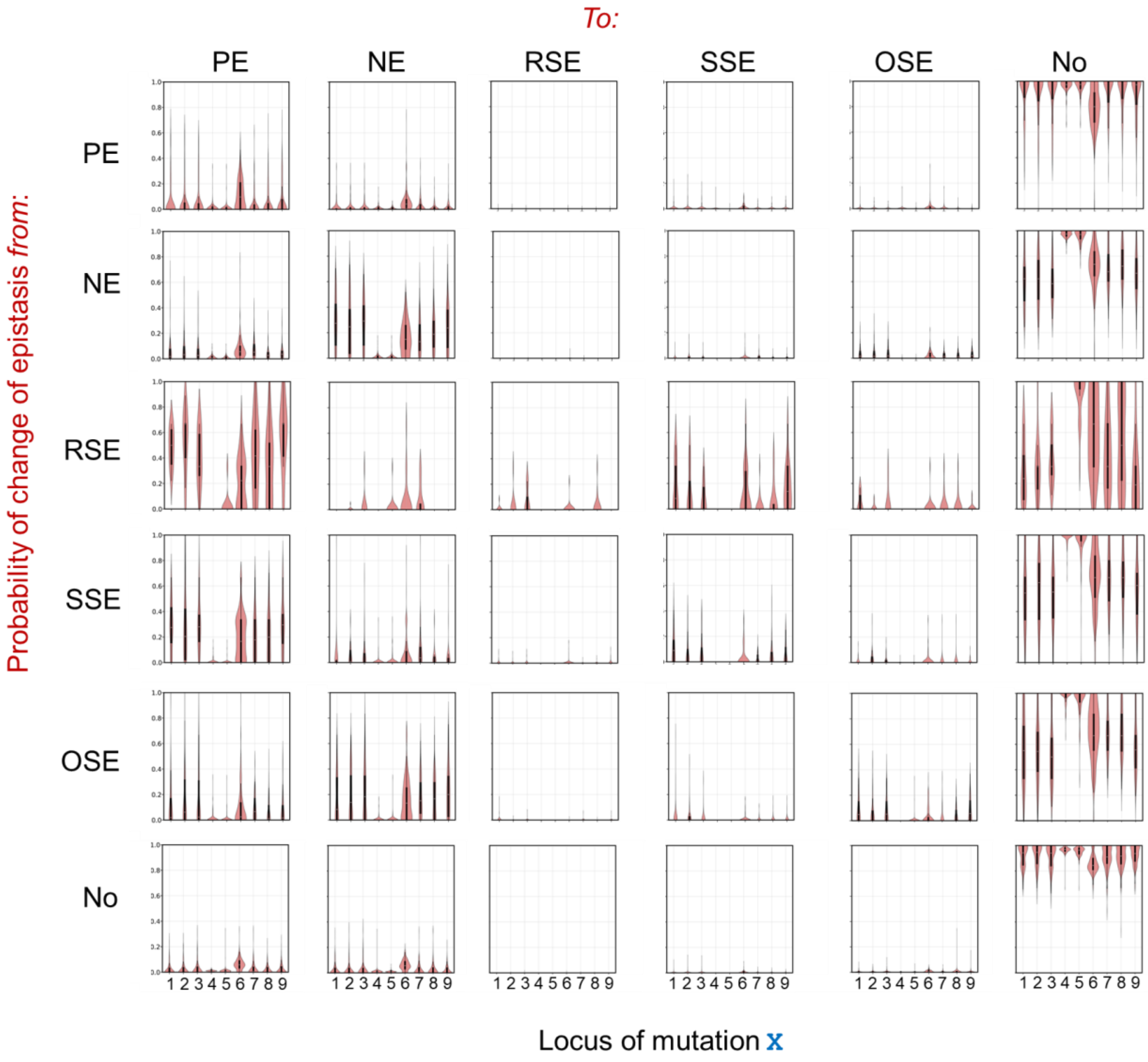


**Figure S9. Change of epistasis because of synonymous mutations as a function of locus *X* in high fitness backgrounds.** Diagonal graphs represent the probability of nature of epistasis not changing. PE: Positive Epistasis; NE: Negative Epistasis; RSE: Reciprocal Sign Epistasis; SSE: Single Sign Epistasis; OSE: Other Sign Epistasis; No: No Epistasis. The probability for change of epistasis was calculated as described in Figure 2A in the main text.

#### Supplement Figure S10.

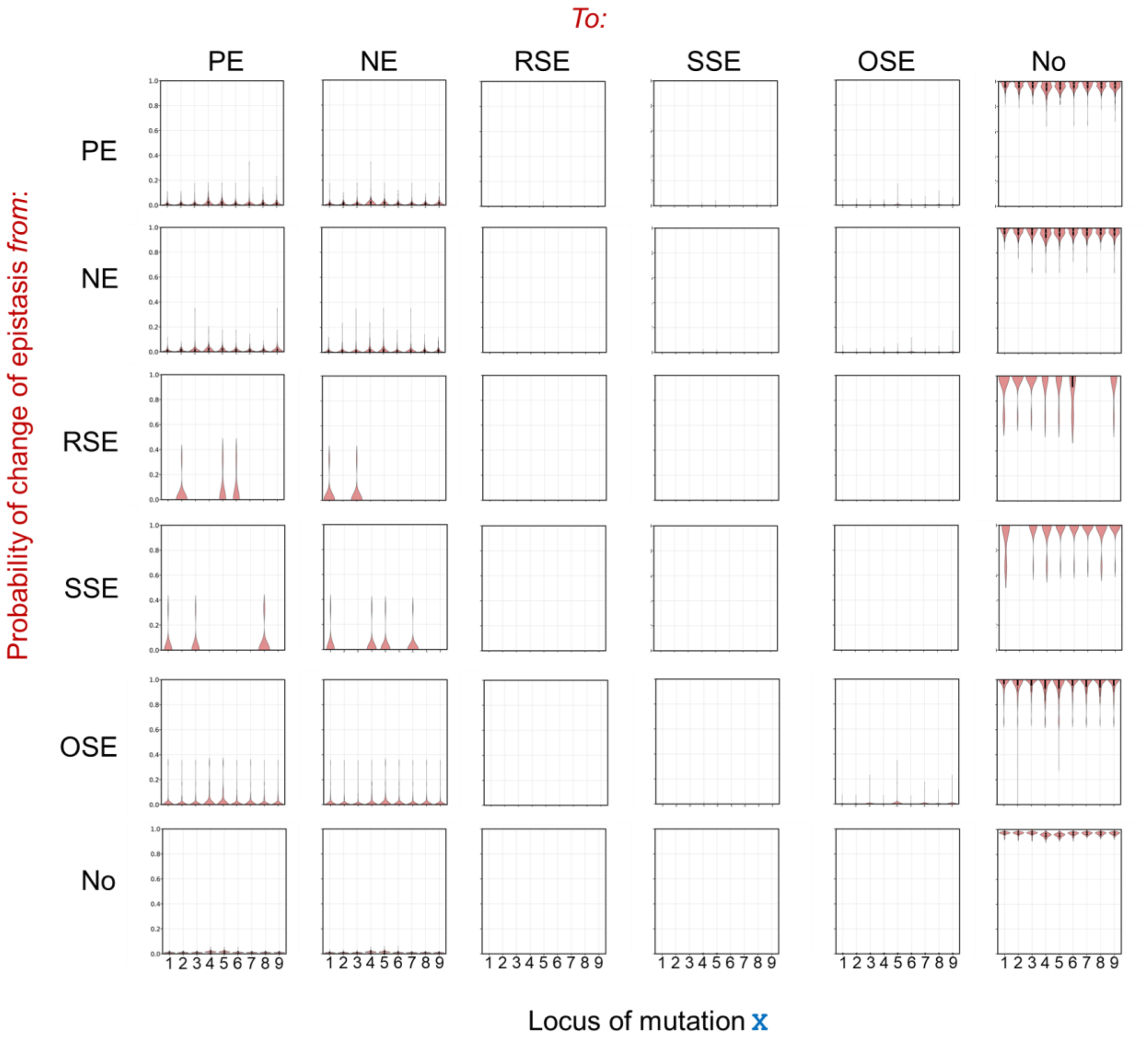


**Figure S10. Change of epistasis because of synonymous mutations as a function of locus *X* in low fitness backgrounds.** Diagonal graphs represent the probability of nature of epistasis not changing. PE: Positive Epistasis; NE: Negative Epistasis; RSE: Reciprocal Sign Epistasis; SSE: Single Sign Epistasis; OSE: Other Sign Epistasis; No: No Epistasis. The probability for change of epistasis was calculated as described in Figure 2A in the main text.

### Supplement S11.

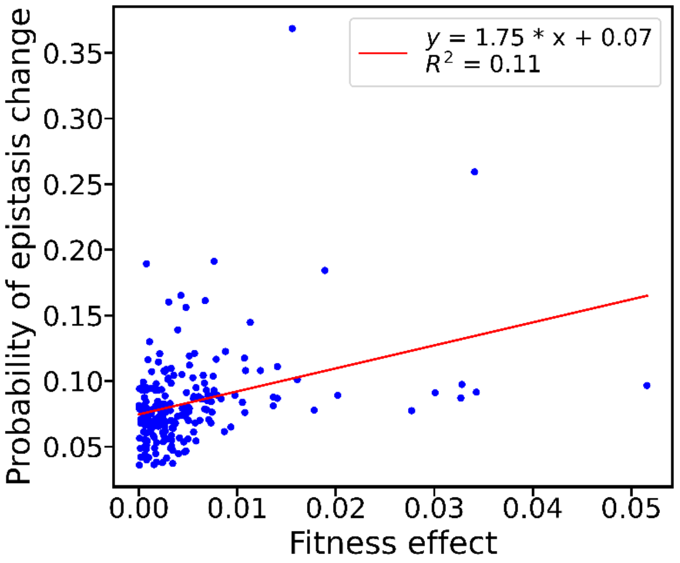


**Figure S11. Relationship between mean fitness effect of synonymous mutations and their influence on epistasis.** Each point represents a synonymous mutation, with the x-axis showing its mean fitness effect and the y-axis showing the percentage of pairwise contexts in which it changes the type of epistasis. No correlation is observed between these two quantities ($R^2 = 0.11$). For clarity, only half of the possible mutational orientations (X → Y) are shown; including the reverse orientations (Y → X) yields a symmetric distribution about the x = 0 axis.

#### Supplement S12.

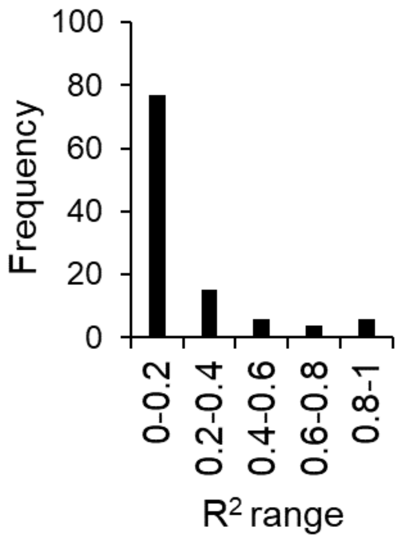


**Figure S12.** A majority (77 in number) mutations show a statistically insignificant correlation ($R^2$ < 0.2) between background fitness and mutational effect.

### Supplement Figure S13.

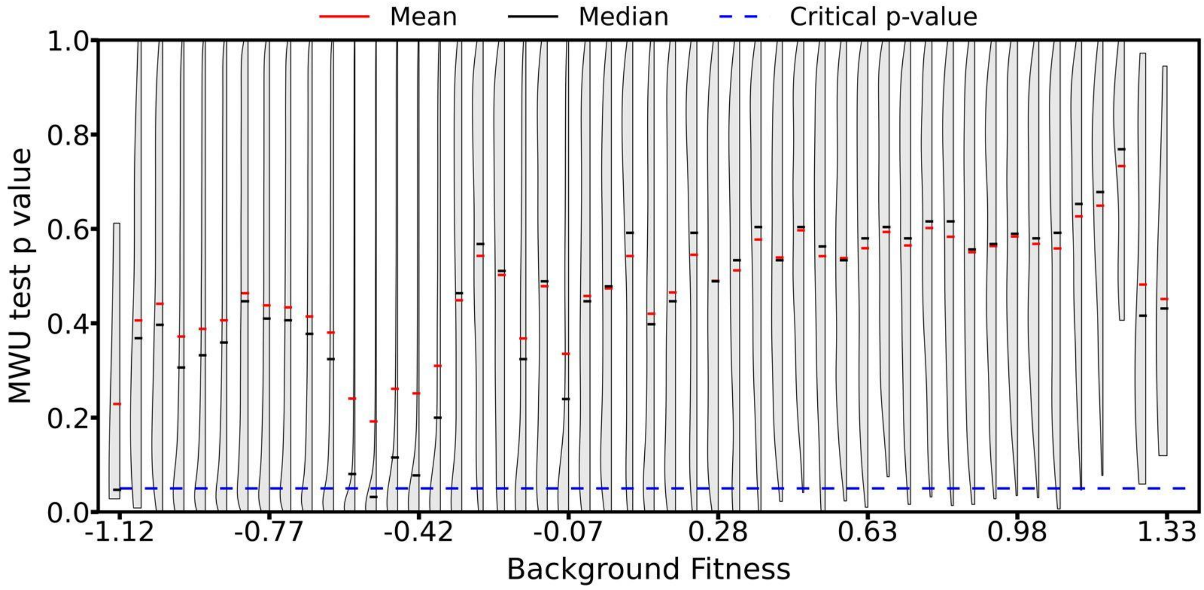


**Figure S13.** Distribution of p-values from an MWU-test to determine statistical similarity between the DFEs of variants with similar fitness. In each band of 0.05 fitness, 90% of the genomes were used to construct the Phenotypic DFE. The DFE of the remaining 10% genomes was compared with the Phenotypic DFE, and the p-value obtained. The distribution of p-value for each background fitness is shown in the Figure. As background fitness increases, Phenotypic DFE becomes a statistically accurate predictor of DFE.

#### Supplement Table 1.

**Table S1. Statistics of the patterns of epistasis on the *folA* landscape.**

| Epistasis | High fitness backgrounds | | Low fitness backgrounds | |
|---|---|---|---|---|
| | **Mean** | **Median** | **Mean** | **Median** |
| **Positive** | 24.573% | 20.970% | 1.654% | 1.040% |
| **Negative** | 13.250% | 10.955% | 2.182% | 1.120% |
| **Reciprocal Sign** | 0.565% | 0.000% | 0.003% | 0.000% |
| **Single Sign** | 1.503% | 0.388% | 0.012% | 0.006% |
| **Other Sign** | 3.475% | 1.656% | 0.266% | 0.119% |
| **No Epistasis** | 55.032% | 56.829% | 95.153% | 96.776% |

#### Supplement Table 2.

**Table S2. p-value from MWU test between distributions for different loci in Figure 2B.**
(distributions which are statistically insignificantly different from each other are in red)

| | | X | | | | | | | | |
|---|---|---|---|---|---|---|---|---|---|---|
| | | 1 | 2 | 3 | 4 | 5 | 6 | 7 | 8 | 9 |
| X | 1 | 1 | | | | | | | | |
| | 2 | 0.2670 | 1 | | | | | | | |
| | 3 | 0 | 0 | 1 | | | | | | |
| | 4 | 0 | 0 | 0 | 1 | | | | | |
| | 5 | 0 | 0 | 0 | 0.0349 | 1 | | | | |
| | 6 | 0.2561 | 0.0218 | 0 | 0 | 0 | 1 | | | |
| | 7 | 0 | 0 | 0 | 1.00E-10 | 0 | 0 | 1 | | |
| | 8 | 0 | 0 | 0 | 5.10E-07 | 0 | 0 | 0.1414 | 1 | |
| | 9 | 0 | 0 | 6.89E-08 | 0 | 0 | 0 | 0 | 0 | 1 |

#### Supplement Table 3.

**Table S3. p-value from MWU test between distributions for different loci in Figure 2C.**
(distributions which are statistically insignificantly different from each other are in red)

| | | X | | | | | | | | |
|---|---|---|---|---|---|---|---|---|---|---|
| | | **1** | **2** | **3** | **4** | **5** | **6** | **7** | **8** | **9** |
| **X** | **1** | 1 | | | | | | | | |
| | **2** | 0.71022 | 1 | | | | | | | |
| | **3** | 0.01865 | 0.04865 | 1 | | | | | | |
| | **4** | 0 | 0 | 0 | 1 | | | | | |
| | **5** | 0 | 0 | 0 | 0.36042 | 1 | | | | |
| | **6** | 0 | 2.00E-10 | 1.14E-05 | 0 | 0 | 1 | | | |
| | **7** | 2.73E-06 | 1.44E-05 | 0.01698 | 0 | 0 | 0.06021 | 1 | | |
| | **8** | 0 | 0 | 0 | 0 | 0 | 0.00676 | 5.00E-06 | 1 | |
| | **9** | 1.00E-09 | 8.50E-09 | 0.00015 | 0 | 0 | 0.54356 | 0.19416 | 0.00112 | 1 |

#### Supplement Table 4.

**Table S4.** Patterns of global epistasis for all 108 mutations in the DHFR landscape.

| Mutation | Position | Linear slope | Intercept | Pivot | $R^2$ |
|---|---|---|---|---|---|
| T -> A | 9 | -0.204 | -0.142 | -0.696 | 0.152 |
| G -> A | 9 | -0.218 | -0.156 | -0.716 | 0.218 |
| C -> A | 9 | -0.242 | -0.169 | -0.698 | 0.206 |
| A -> T | 9 | -0.080 | -0.045 | -0.556 | 0.020 |
| G -> T | 9 | -0.144 | -0.099 | -0.691 | 0.108 |
| C -> T | 9 | -0.091 | -0.062 | -0.681 | 0.078 |
| A -> G | 9 | -3e-6 | 0.015 | 446.666 | 0.000 |
| T -> G | 9 | -0.052 | -0.028 | -0.535 | 0.013 |
| C -> G | 9 | -0.090 | -0.055 | -0.612 | 0.040 |
| A -> C | 9 | -0.052 | -0.023 | -0.448 | 0.008 |
| T -> C | 9 | -0.017 | -0.008 | -0.494 | 0.002 |
| G -> C | 9 | -0.110 | -0.075 | -0.679 | 0.061 |
| T -> A | 8 | -0.553 | -0.401 | -0.725 | 0.596 |
| G -> A | 8 | -0.329 | -0.234 | -0.711 | 0.279 |
| C -> A | 8 | -0.506 | -0.350 | -0.692 | 0.374 |
| A -> T | 8 | 0.096 | 0.124 | -1.293 | 0.007 |
| G -> T | 8 | 0.074 | 0.092 | -1.233 | 0.009 |
| C -> T | 8 | 0.026 | 0.062 | -2.371 | 0.001 |
| A -> G | 8 | -0.081 | -0.038 | -0.472 | 0.012 |
| T -> G | 8 | -0.400 | -0.290 | -0.724 | 0.446 |
| C -> G | 8 | -0.205 | -0.134 | -0.654 | 0.126 |
| A -> C | 8 | -0.265 | -0.164 | -0.620 | 0.069 |
| T -> C | 8 | -0.381 | -0.279 | -0.734 | 0.397 |
| G -> C | 8 | -0.138 | -0.093 | -0.672 | 0.053 |
| T -> A | 7 | -0.125 | -0.073 | -0.584 | 0.054 |
| G -> A | 7 | -0.114 | -0.065 | -0.569 | 0.060 |
| C -> A | 7 | -0.230 | -0.148 | -0.643 | 0.106 |
| A -> T | 7 | -0.159 | -0.112 | -0.706 | 0.091 |

| | | | | | |
|---|---|---|---|---|---|
| G -> T | 7 | -0.206 | -0.135 | -0.654 | 0.117 |
| C -> T | 7 | -0.242 | -0.165 | -0.682 | 0.121 |
| A -> G | 7 | -0.103 | -0.077 | -0.748 | 0.049 |
| T -> G | 7 | -0.163 | -0.108 | -0.664 | 0.070 |
| C -> G | 7 | -0.265 | -0.181 | -0.682 | 0.121 |
| A -> C | 7 | -0.259 | -0.170 | -0.656 | 0.140 |
| T -> C | 7 | -0.241 | -0.151 | -0.627 | 0.120 |
| G -> C | 7 | -0.302 | -0.191 | -0.631 | 0.165 |
| T -> A | 6 | -0.256 | -0.189 | -0.740 | 0.212 |
| G -> A | 6 | -0.040 | -0.025 | -0.623 | 0.012 |
| C -> A | 6 | -0.257 | -0.189 | -0.737 | 0.206 |
| A -> T | 6 | -0.066 | -0.019 | -0.282 | 0.011 |
| G -> T | 6 | -0.050 | -0.007 | -0.130 | 0.006 |
| C -> T | 6 | -0.054 | -0.034 | -0.628 | 0.025 |
| A -> G | 6 | -0.090 | -0.062 | -0.686 | 0.063 |
| T -> G | 6 | -0.283 | -0.209 | -0.737 | 0.250 |
| C -> G | 6 | -0.283 | -0.208 | -0.735 | 0.242 |
| A -> C | 6 | -0.079 | -0.028 | -0.352 | 0.016 |
| T -> C | 6 | -0.064 | -0.042 | -0.652 | 0.035 |
| G -> C | 6 | -0.062 | -0.015 | -0.241 | 0.009 |
| T -> A | 5 | -0.417 | -0.047 | -0.114 | 0.003 |
| G -> A | 5 | -1.174 | -0.593 | -0.505 | 0.072 |
| C -> A | 5 | -0.900 | -0.400 | -0.444 | 0.019 |
| A -> T | 5 | -0.990 | -0.723 | -0.730 | 0.984 |
| G -> T | 5 | -0.999 | -0.726 | -0.727 | 0.771 |
| C -> T | 5 | -0.994 | -0.723 | -0.727 | 0.586 |
| A -> G | 5 | -1.010 | -0.704 | -0.698 | 0.948 |
| T -> G | 5 | -0.995 | -0.696 | -0.700 | 0.226 |
| C -> G | 5 | -0.802 | -0.558 | -0.695 | 0.217 |
| A -> C | 5 | -0.998 | -0.718 | -0.720 | 0.977 |
| T -> C | 5 | -0.991 | -0.713 | -0.719 | 0.406 |
| G -> C | 5 | -0.917 | -0.661 | -0.721 | 0.669 |

| | | | | | |
|---|---|---|---|---|---|
| **T -> A** | **4** | -0.904 | -0.655 | -0.725 | 0.721 |
| **G -> A** | **4** | -0.998 | -0.722 | -0.723 | 0.983 |
| **C -> A** | **4** | -0.988 | -0.714 | -0.723 | 0.547 |
| **A -> T** | **4** | -0.706 | -0.489 | -0.693 | 0.143 |
| **G -> T** | **4** | -1.020 | -0.711 | -0.697 | 0.952 |
| **C -> T** | **4** | -0.993 | -0.697 | -0.702 | 0.285 |
| **A -> G** | **4** | -0.875 | -0.377 | -0.431 | 0.013 |
| **T -> G** | **4** | -1.381 | -0.734 | -0.532 | 0.092 |
| **C -> G** | **4** | -0.302 | 0.038 | 0.126 | 0.002 |
| **A -> C** | **4** | -0.985 | -0.713 | -0.724 | 0.439 |
| **T -> C** | **4** | -0.997 | -0.722 | -0.724 | 0.711 |
| **G -> C** | **4** | -0.985 | -0.717 | -0.728 | 0.979 |
| **T -> A** | **3** | -0.202 | -0.145 | -0.716 | 0.172 |
| **G -> A** | **3** | -0.124 | -0.089 | -0.718 | 0.104 |
| **C -> A** | **3** | -0.202 | -0.143 | -0.709 | 0.164 |
| **A -> T** | **3** | -0.043 | -0.015 | -0.344 | 0.006 |
| **G -> T** | **3** | -0.048 | -0.026 | -0.534 | 0.013 |
| **C -> T** | **3** | -0.049 | -0.030 | -0.614 | 0.021 |
| **A -> G** | **3** | -0.028 | -0.011 | -0.379 | 0.005 |
| **T -> G** | **3** | -0.119 | -0.083 | -0.698 | 0.087 |
| **C -> G** | **3** | -0.120 | -0.082 | -0.683 | 0.081 |
| **A -> C** | **3** | -0.057 | -0.026 | -0.453 | 0.011 |
| **T -> C** | **3** | -0.063 | -0.043 | -0.677 | 0.036 |
| **G -> C** | **3** | -0.063 | -0.037 | -0.593 | 0.021 |
| **T -> A** | **2** | -0.156 | -0.107 | -0.684 | 0.104 |
| **G -> A** | **2** | -0.091 | -0.062 | -0.684 | 0.060 |
| **C -> A** | **2** | -0.070 | -0.048 | -0.683 | 0.022 |
| **A -> T** | **2** | -0.086 | -0.051 | -0.598 | 0.029 |
| **G -> T** | **2** | -0.104 | -0.066 | -0.632 | 0.045 |
| **C -> T** | **2** | -0.051 | -0.030 | -0.595 | 0.009 |
| **A -> G** | **2** | -0.049 | -0.029 | -0.600 | 0.017 |
| **T -> G** | **2** | -0.135 | -0.090 | -0.668 | 0.077 |

| | | | | | |
|---|---|---|---|---|---|
| **C -> G** | **2** | -0.062 | -0.040 | -0.642 | 0.015 |
| **A -> C** | **2** | -0.144 | -0.092 | -0.643 | 0.099 |
| **T -> C** | **2** | -0.194 | -0.129 | -0.667 | 0.158 |
| **G -> C** | **2** | -0.175 | -0.115 | -0.658 | 0.133 |
| **T -> A** | **1** | 0.020 | 0.040 | -2.024 | 0.001 |
| **G -> A** | **1** | 0.080 | 0.067 | -0.842 | 0.026 |
| **C -> A** | **1** | -0.013 | 0.004 | 0.281 | 0.001 |
| **A -> T** | **1** | -0.318 | -0.231 | -0.725 | 0.332 |
| **G -> T** | **1** | -0.190 | -0.137 | -0.720 | 0.114 |
| **C -> T** | **1** | -0.269 | -0.191 | -0.707 | 0.241 |
| **A -> G** | **1** | -0.230 | -0.162 | -0.705 | 0.306 |
| **T -> G** | **1** | -0.135 | -0.078 | -0.577 | 0.054 |
| **C -> G** | **1** | -0.168 | -0.112 | -0.668 | 0.188 |
| **A -> C** | **1** | -0.136 | -0.099 | -0.732 | 0.125 |
| **T -> C** | **1** | -0.042 | -0.014 | -0.321 | 0.005 |
| **G -> C** | **1** | 0.022 | 0.017 | -0.770 | 0.003 |

### Supplement References.


1 Papkou, A., Garcia-Pastor, L., Escudero, J. A. & Wagner, A. A rugged yet easily navigable fitness landscape. *Science* **382**, eadh3860, doi:10.1126/science.adh3860 (2023).

2 Kauffman, S. & Levin, S. Towards a general theory of adaptive walks on rugged landscapes. *J Theor Biol* **128**, 11-45, doi:10.1016/s0022-5193(87)80029-2 (1987).

3 Weinreich, D. M., Delaney, N. F., Depristo, M. A. & Hartl, D. L. Darwinian evolution can follow only very few mutational paths to fitter proteins. *Science* **312**, 111-114, doi:10.1126/science.1123539 (2006).
